# *Pellaea zygophylla*, a new combination for a distinctive & well-known but neglected fern

**DOI:** 10.1101/2021.03.25.437027

**Authors:** Patrick J. Alexander

**Author notes:** /.

## Abstract

*Pellaea ovata* is a widespread species, sexual diploid in Texas &northeastern Mexico but an apogamous triploid in northwestern Mexico, south to northern Argentina, &on Hispaniola. The type belongs to the southern, apogamous triploid form. Although these two forms have been discussed repeatedly in the literature, morphological distinctions between them have been overlooked and they have not been recognized taxonomically. However, they are distinct. *Pellaea ovata* s.s. has puberulent rachides & costae; pinnae usually 2-pinnate with a well-defined main axis &pinnules borne singly; fertile pinnules ovate, cordate basally &rounded apically. The sexual diploid form has rachides &costae glabrous; pinnae pseudo-dichotomously branched &pinnules usually paired; fertile pinnules narrowly rounded-trapeziform, obliquely truncate to cordate basally &truncate apically. Riddell named the sexual diploid form *Pteris zygophylla*, from which I give it the new combination *Pellaea zygophylla*.

---

I first encountered *Pellaea ovata* in the greenhouse of Dr. Gerald J. Gastony at Indiana University in late 2002. In 2003 &2004 I noticed two distinct morphologies among his plants. One form had glabrous rachides &almost dichotomous branching in the pinnae, the other puberulent rachides &pinnae more straightforwardly pinnate. I began to suspect that these may be separate species, a suspicion that lingered in the back of my mind over the following years. After seeing live plants in the field in central Texas in August 2006 &checking the literature, I learned that these forms are correlated with ploidy &reproductive mode. The Texas plants belonged to the glabrous, dichotomous form, and Texas plants were reported to be sexual &diploid (Tryon 1957, Tryon &Britton 1958, Tryon 1968, Tryon 1972). The pubescent, pinnate form must correspond with the reported apomictic triploids, then. Consulting Mickel &Smith (2004) around the same time it became clear to me that the two forms of *Pellaea ovata* cannot be understood separately from *Pellaea oaxacana*. Segregated from *Pellaea ovata* by Mickel &Beitel (1988), *Pellaea oaxacana* resembles the Texas plants in having rachides &costae glabrous or nearly so. The rachides &cosate are less flexuous than in either form of *Pellaea ovata* and the pinnules are not paired. Flexuosity varies in all of these taxa, however, and the distinction between *Pellaea oaxacana* and *Pellaea ovata* s.l. is not entirely clear. Recently, this taxonomic question regained my attention while reviewing observations on iNaturalist. I found an image of the type of *Pteris zygophylla* at Gray Herbarium (gh 339038) and realized that a name had been published for the sexual diploid form of *Pellaea ovata*. Riddell’s name needs only to be revived and combined in *Pellaea*. In this paper I do so, provide morphological descriptions &distribution maps for this species along with *Pellaea oaxacana* and *Pellaea ovata*, and discuss some of the remaining uncertainties surrounding these species.

John Riddell (1853) published new names for plants collected in Louisiana and Texas, the Texas plants having been collected in 1839. These names came primarily from an unpublished flora of Louisiana that he sent, with specimens and illustrations, to the Smithsonian. This unpublished manuscript and its associated materials were sent to Gray Herbarium, where they now reside. Riddell’s 1853 names have generally been overlooked. They are not mentioned in any work on *Pellaea* until Wilbur &Whitson (2005) brought attention to Riddell’s seven fern basionyms, *Pteris zygophylla* among them. Wilbur &Whitson republished the description of *Pteris zygophylla*, found the illustration at gh, and indicated that the name is a synonym of *Pellaea ovata*. Although Riddell does not mention *Pellaea ovata*, his description includes most of the features distinguishing his *Pteris zygophylla* from that species. Like Wilbur &Whitson (2005), I provide it in full:

Pteris zygophylla. *Frond* glabrous, supra-decompound, outline triangular lanceolate; *subdivisions* of the stipe alternate, petiolate, divaricate; *pinnules* mostly in pairs (zygophyllous), trapeziform, sub-ovate, obliquely cordate at base; apex truncate, (about half inch long by one third or one fourth inch broad) *veins* immersed in the substance of the pinnule; *veinlets* once or twice forked near the lateral margin, where they bear the *sporangia*, which form a marginal spore extending the whole length of each pinnule on each side, more or less covered by the reflected membranaceous margin of the pinnule; *stipe* yellowish brown, smooth above, chaffy near the roots, sub-scandent; about two feet high. Grows among granite rocks in the mountains of the Camanche country, Texas. (Oct. 1839.) Natural order Filices.

Within this description, the following features are especially relevant: stipe &frond glabrous; pinnules mostly in pairs, trapeziform, apex truncate. *Pellaea ovata* has the stipe and frond (rachis, costae, stalks of the pinnules, occasionally surfaces of the pinnules) puberulent; pinnules mostly single, ovate, apex rounded. Subsequent treatments have generally neglected these features along with Riddell’s publication, highlighting the sexual diploid and apogamous triploid forms of *Pellaea ovata* but rarely mentioning morphological distinctions beyond cell size or spore features directly related to ploidy &reproduction (Tryon 1957; Tryon 1968; Tryon &Britton 1958; Tryon 1972; Windham 1993; Wilbur &Whitson 2005). Tryon (1957) provides a noteworthy exception. Although she describes *Pellaea ovata* as “relatively uniform throughout most of its range” and attributes this to apogamy in most of the range, in later discussion of *Pellaea sagittata* (her *Pellaea sagittata* var. *sagittata*) she writes:

The presence of pubescence, particularly on the rachises, appears to be correlated with the apogamous condition. In *P. ovata* and *P. andromedaefolia*, as well as in this variety, it is a convenient clue for detecting specimens with 32 spored sporangia and apparently apogamous.

In developing the morphological descriptions and distribution maps below, I have relied heavily on digital records. I reviewed images of a total of 837 accessions of *Pellaea ovata* sensu lato &*Pellaea oaxacana*. This includes including 419 herbarium specimen images, accessed from the following portals:

> PteridoPortal (pteridoportal.org),
>
> IBData (ibdata.ib.unam.mx),
>
> SEINet (swbiodiversity.org),
>
> CCH2 (cch2.org).

Herbarium specimens are cited below; only those of which I saw an image are cited. I also reviewed 418 photographic observations on iNaturalist (inaturalist.org). A data file citing both herbarium specimens and photographic observations is available on Data Dryad at https://doi.org/10.5061/dryad.0vt4b8gzg. I also reviewed related taxa, especially *Pellaea cordifolia* (Sessé &Moc.) A.R.Sm. &*Pellaea sagittata* (Cav.) Link, but do not include extensive citations for these. The availability of large numbers of specimen images &photographic observations makes it easy to review large numbers of plants across the globe, although the high volume of observations is counteracted by reduced information per observation. Photographs &specimen images are never as good as viewing a plant under a dissecting microscope, and of course it is impossible to count spores. Luckily, most of the features relevant to *Pellaea ovata* s.l. are macroscopic &discernible in good images of live plants or specimens, although pubescence is not always apparent. I have used the descriptions of Tryon (1957), Windham (1993), Mickel &Smith (2004), &Velázquez-Montes (2018) as a starting point in developing descriptions, and as the primary source for features like rhizome scales that are difficult to discern in specimen images. Descriptions of frond features apply to fully developed, fertile leaves on mature plants. Fronds are often scalier or more pubescent as they unfurl, and glabrescent with age. Young or small plants may also differ in their morphology, tending especially to have straighter rachides &costae. Mature plants, especially of *Pellaea ovata* s.s., sometimes produce anomalously large pinnules on early-season, sterile leaves, or on sterile pinnae near the base of distally fertile leaves.

> Pellaea zygophylla (Riddell) P.J.Alexander, comb. nov.
>
> *Pteris zygophylla* Riddell, New Orleans Med. Surg. J. 9: 616 (1853). Type: *Riddell s*.*n*., Oct 1839, Comanche country, Texas (gh 339038!, NY 3496495!).
>
> Figure 1 (a, b), Figures 2–6.

Rhizomes creeping, slender, 2–4 mm in diameter; *scales* loosely appressed, lanceolate, 2–3 × 0.3–0.8 mm, bicolorous, centers black, dull or weakly lustrous, with thin, brown, erose-dentate margins. Leaves 20–80(–120) × (5–)7–15(–20) cm, ascending to sprawling, sometimes subscandent; *stipe* 0.8–1.1 times as long as the blade, rounded or flattened adaxially, scaly for 1–3 cm at the base, basalmost scales persistent, dense, like those of the rhizome, more distal scales gradually deciduous, sparse, pale, and linear; stipe and rachis tan to reddish-brown, turning very pale gray with age; *rachis* weakly to strongly flexuous, rarely straight, glabrous; *blade* lanceolate, usually 3-pinnate, occasionally 2-pinnate, with (4–)6–10(–15) pairs of pinnae, alternate or (rarely) subopposite; distalmost 2–5 pinnules borne singly on the rachis. Pinnae reflexed to slightly ascending, the larger pinnae typically 2-pinnate, with (2–)4–12(–20) pinnules, many of them distinctly paired, branching of the pinnae appearing almost dichotomous and the central axis not readily apparent, each node a broad **Y** or **T** with the stem towards the rachis, somewhat oblique to equilateral; *costae* strongly flexuous, base strongly to weakly reflexed, glabrous; *stalks* of the pinnules 1–6(–10) mm, usually with a few translucent multicellular trichomes 0.1–0.3 mm long near the bases of the pinnules, sometimes sparsely puberulent about half their length; costae &stalks the same color as the rachis, often darker immediately at the base of a pinnule. Pinnules trapeziform to rounded-trapeziform, occasionally broadly lanceolate (especially on smaller leaves), (7–)9–25(–30) × (3–)4–12(–16) mm, 2–2.5 times longer than wide, coriaceous, glabrous, veins indistinct; *base* truncate or widely cordate, oblique or (rarely) equilateral; *apex* truncate or (rarely) rounded-acute, almost always with a pronounced gap between the sori; *sori* not visible adaxially, false indusia 0.3–0.7 mm wide, revolute, entire, thinning and becoming pale at the margin but otherwise little differentiated from the rest of the pinnule.

64 spores per sporangium; plants sexual, diploid. Tryon &Britton (1958) and Tryon (1968) report sexual diploids, 2n = 58, from central Texas (*Tryon &Tryon 5029, Tryon &Tryon 5524*; not seen). Tryon also counted spores, finding only 32-spored plants in Texas &northeastern Mexico, but does not cite specimens for the counts.

Central Texas (Palo Pinto County) and south, mostly along the east side of the Sierra Madre Orientál, to the state of Morelos. Map 1 &Map 2. Mostly on semiarid limestone.

Mexico. Coahuila: *Encina &al. 1634* (mexu 1372693); *Palmer s*.*n*. (yu 20021); *Pinkava &Reeves R-4329* (huap 27834, mexu 1403971); *Wynd &Mueller 318* (us 1639759). Nuevo LeÓn: *Briones 1883* (brit 432135); *Copeland s*.*n*. (mich 1208316); *Dorr &al. 2575* (uc 1513612); *Estrada 16202* (brit 432136); *Fryxell &Kirkpatrick 2469* (vt 286371); *Gastony &Yatskievych 86-24* (ind 3412); *Hinton &Hinton 21460* (mo 3605676); *Hinton 21140* (mo 3605321); *Kimber s*.*n*. (ph 737306); *Knobloch 2017* (msc 267092); *McCulloch 76-71-Mc* (msc 267090); *Palmer s*.*n*. (yu 20022); *Pennell 16954* (huap 27834, mexu 1403971); *Rodríguez 88* (mexu 821157); *Storer 68* (mich 1208287). San Luis PotosÍ: *Gastony &Yatskievych 86-27* (ind 3409); *Pringle s*.*n*. (huap 27834, mexu 1403971). Tamaulipas: *Bartlett 10183* (mich 1208222); *Bartlett 10313* (mexu 88785, mich 1208286, us 1490578); *Bartlett 10658* (mich 1208223); *Bartlett 10707* (mich 1208291); *Bartlett 10802* (mich 1208219, us 1490603); *Briones 1234* (mexu 844575); *Knobloch 2245* (f 633209, msc 267094); *Runyon 717* (brit 432123); *Walker &Baker 2088* (wis 113330); *Windham &al. 500* (ut 99958); *Yatskievych &Gastony 86-44* (ind 136927).

U.S.A. T exas : *Atha 11729* (ny 1745374); *Barkley &al. 47252* (ph 737502); *Blassingame 2811* (hpc 16817); *Buckley s*.*n*. (ny 3496505); *Carloyne 53* (hpc 16816); *Correll 13464* (ny 3496490); *Ertter 4904* (ny 3496486); *Ferriss s*.*n*. (ph 737505 &737506); *Gerault 6* (hpc 25769); *Gowdy 53* (hpc 25765); *Gowdy 7* (hpc 25771); *Hill 8658* (vt 286372); *Lindheimer 1280* (ny 3496496, ph 737498); *Lott &Rankin 4644* (tenn 4559); *Mohr s*.*n*. (missa 263); *Palmer 1428* (ny 3496491, ph 737501); *Parks s*.*n*. (ph 737510); *Pilsbry s*.*n*. (ph 737499); *Pilsbry s*.*n*. (ph 737504); *Plank s*.*n*. (ny 3496508); *Pray 1728* (ny 3496483); *Ragsdale 116* (hpc 25756); *Reverchon 1628* (ny 3496485, 3496493, 3496504, &3496506); *Reverchon 79* (ind 3405); *Reverchon s*.*n*. (ny 3496507); *Simpson 187b* (sat 12663); *Stanfield s*.*n*. (ny 3496499 &3496509); *Stanford 4246* (hpc 16891, 16893, &25751); *Tharp &Whitehouse s*.*n*. (ph 737503); *Tharp 47252* (ind 3406); *Thomas 8* (hpc 25757); *Wagner 32* (hpc 25947); *Walters 7* (hpc 25762); *Wherry s*.*n*. (ph 737500); *White 86* (hpc 16807); *Windham &al. 4428* (ut 100004).

## Typification of *Pteris zygophylla*

In addition to illustrations &specimens that Riddell sent to the Smithsonian (now at gh) for his flora of Louisiana, Riddell had sent Texas plants to John Torrey not long after collecting them in 1839. Several are cited by Torrey &Gray (1843). These specimens are now at the New York Botanical Garden &include a sheet of *Pellaea ovata* (ny 3496495). Written on the sheet is “*P. divaricata* Ridd. mss.” The name does not appear in his published work, so Riddell apparently sent an unpublished manuscript with the specimens. *Senecio fragrans* Riddell must also have been in this manuscript, as Torrey &Gray (1843) attribute the name to “Ridd. mss.” but neither is it in Riddell’s published work. A third name that must be from this manuscript, *Melothria coccinea*, is written on a sheet at ny (172476). Riddell later (1853) published this species as *Melothria punctata*. In any case, the name “*P. divaricata*” was never published &thus plays no role in priority or other nomenclatural issues under ICNafp. It does, however, help link the sheet at ny with *Pteris zygophylla*. Riddell apparently decided this was a new species shortly after collecting it. He sent supporting material &manuscripts first to Torrey, later to the Smithsonian. Although he changed the name for his new *Pteris* over time, his “*P. divaricata*” &eventual *Pteris zygophylla* are surely from the same material and intended as names for the same taxon. Both gh &ny sheets, then, are original material for his taxonomic concept, and both are types. The number, 1773, that accompanies the sheet at gh appears to be a plate number for his flora rather than a collection number in the usual sense, so I refer to both as “*Riddell s*.*n*.”

## Discussion

Although the branching of the pinnae is difficult to describe adequately in words, it is very distinctive and allows most specimens of *Pellaea zygophylla* to be identified at a glance. When more than a glance is required, the glabrous rachides &costae of this species distinguish it from *Pellaea ovata*; its flexuous rachides &costae and paired pinnules distinguish it from *Pellaea oaxacana*. Difficulties in identifying *Pellaea zygophylla* are generally limited to incomplete information. Young plants with pinnate or barely 2-pinnate leaves can generally still be distinguished from *Pellaea ovata* by pubescence, but sometimes not from *Pelleaa oaxacana*. Specimens of *Pellaea ovata* consisting of old and tangled leaves, with the branching difficult to discern and much of the pubescence lost to age, can occasionally be difficult to distinguish from *Pellata zygophylla*. Specimens from near La Natividad, Oaxaca (*Mickel &Hellwig 3706*, uc 1494872 & ny 3902407; *Yatskievych &González 85-210*, ind 3413) are the only I have found that appear to be truly intermediate between *Pellaea zygophylla* and another species The branching of the pinnae is reminiscent of *Pellaea zygophylla*, although with few of the pinnules paired. The costae &stalks of the pinnules appear to be too pubescent for *Pellaea zygophylla*, but near or a little past the glabrescent extreme of *Pellaea ovata*; pinnule size and shape appear typical of *Pellaea ovata*. These plants are ±200 miles southeast of the nearest known *Pellaea zygophylla* and I think they are more likely an aberrant form of *Pellaea ovata*.

> *Pellaea ovata* (Desv.) Weath.
>
> Contr. Gray Herb. 114: 34 (1936).
>
> *Pteris ovata* Desv., Mem. Soc. Linn. Paris 6(3): 301 (1827). Type:
>
> *Anonymous, s*.*n*., s.d., Peru (p 586562!).
>
> *Hemionitis ovata* (Desv.) Christenh., Global Fl. 4: 18 (2018).
>
> *Pteris flexuosa* Kaulf. ex Schlecht. &Cam., Linnaea 5(4): 614 (1930). Type: *Schiede s*.*n. “785”*, May 1839, Jalapa, Mexico (hal 137767!, hal 137766!, b 20 0103148!, le 8610!).
>
> *Allosorus flexuosus* (Kaulf. ex Schlecht. &Cam.) Kze., Linnaea 13: 136 (1839).
>
> *Pellaea flexuosa* (Kaulf. ex Schlecht. &Cam.) Link, Fil. Spec. 60 (1841).
>
> *Platyloma flexuosum* (Kaulf. ex Schlecht. &Cam.) J.Sm., Bot. Mag. 72 (Companion): 21 (1846).
>
> Figure 1 (c, d), Figures 7–13.

**Figure 1.**
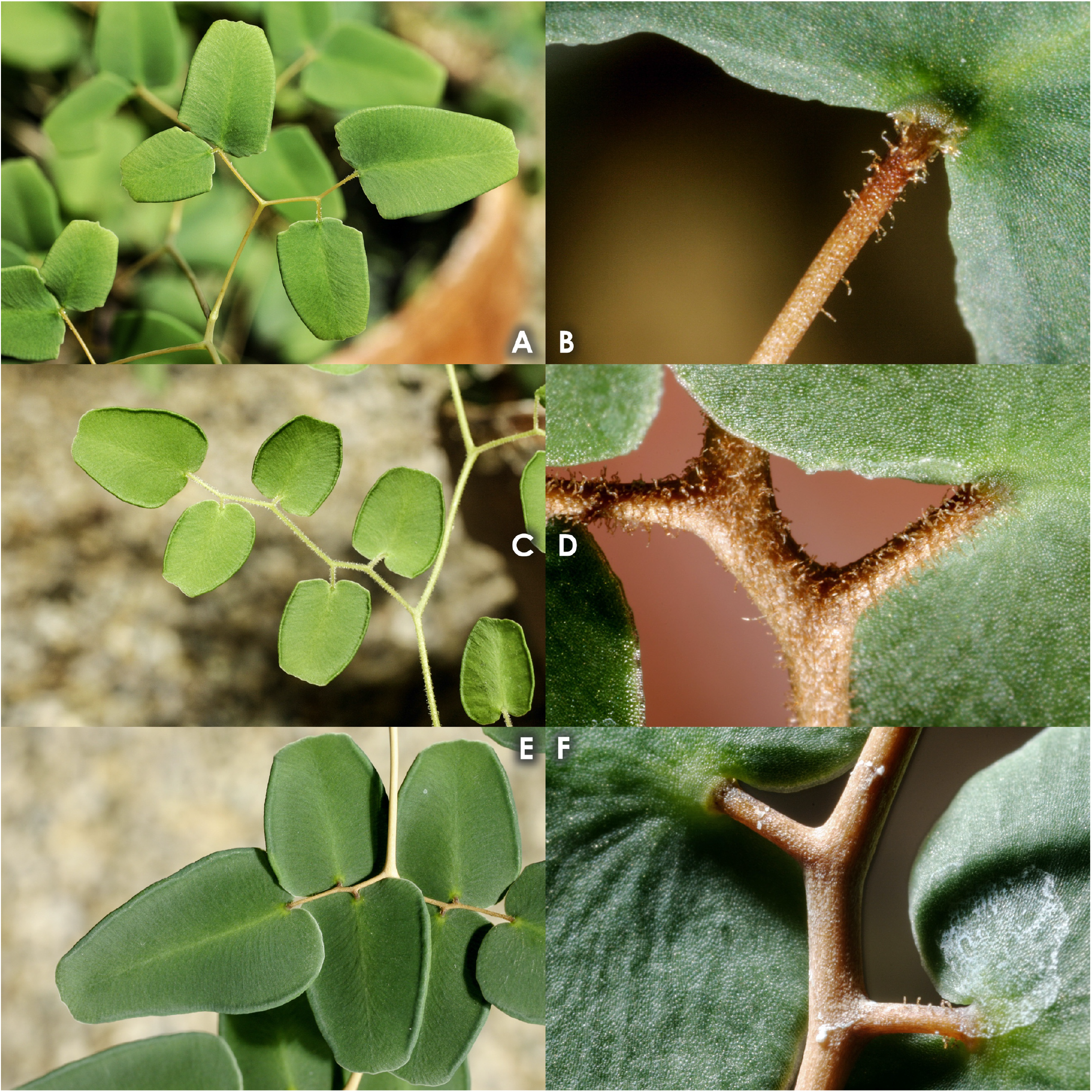
*Pellaea zygophylla* (**A, B**), *Pellaea ovata* (**C, D**), and *Pellaea oaxacana* (**E, F**) in Gastony’s greenhouse, February 2004. Pinnae shown at left, pubescence of the stalks of the pinnules / costae shown at right. Although these plants are surely associated with herbarium specimens, unfortunately I do not have the accession information. This *Pellaea ovata* has pinnule apices at the truncate extreme of variation in the species.

**Figure 2.**
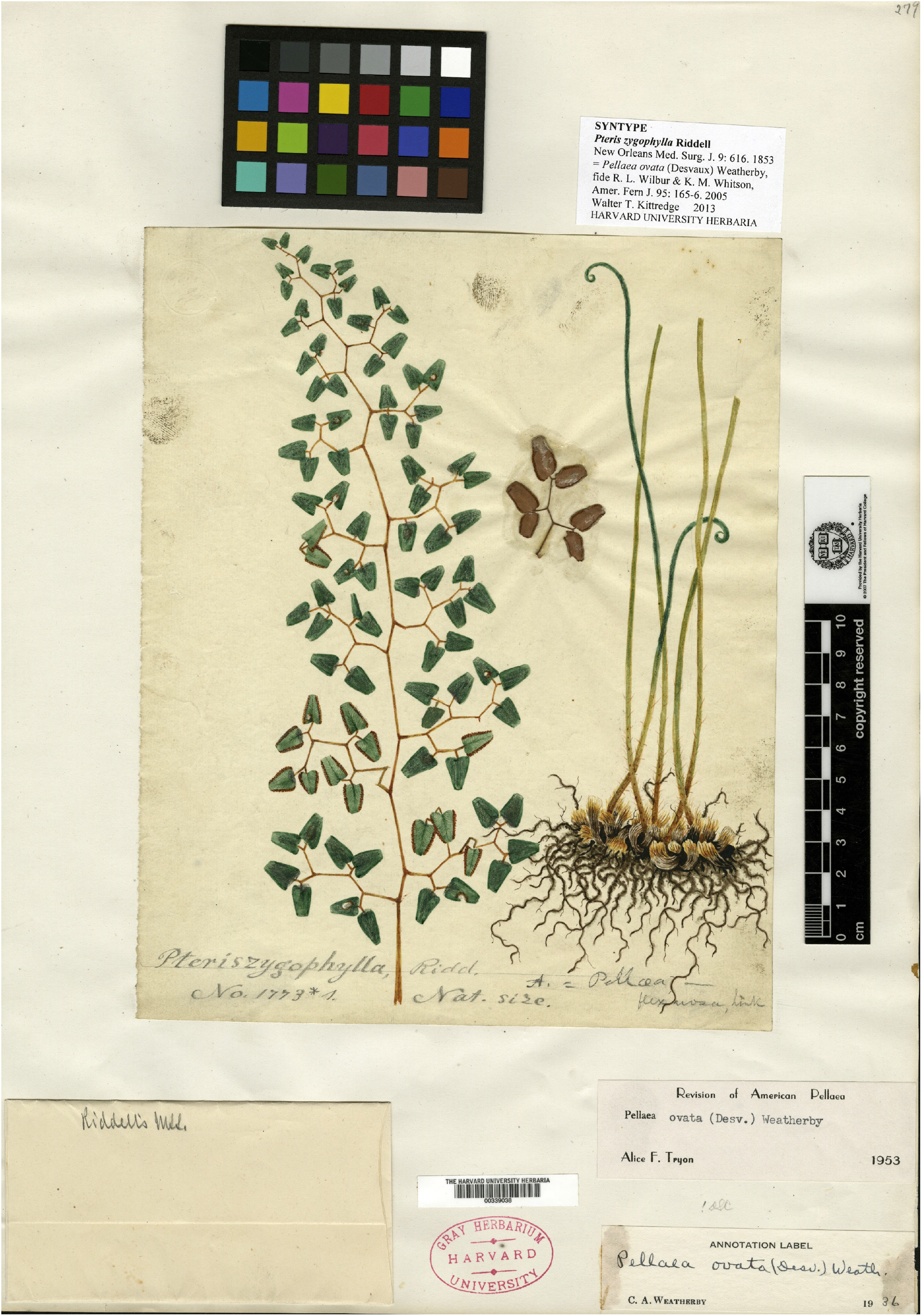
Type of *Pteris zygophylla*: *Riddell s*.*n*., Oct 1839, Comanche country, Texas (gh 339038).

**Figure 3.**
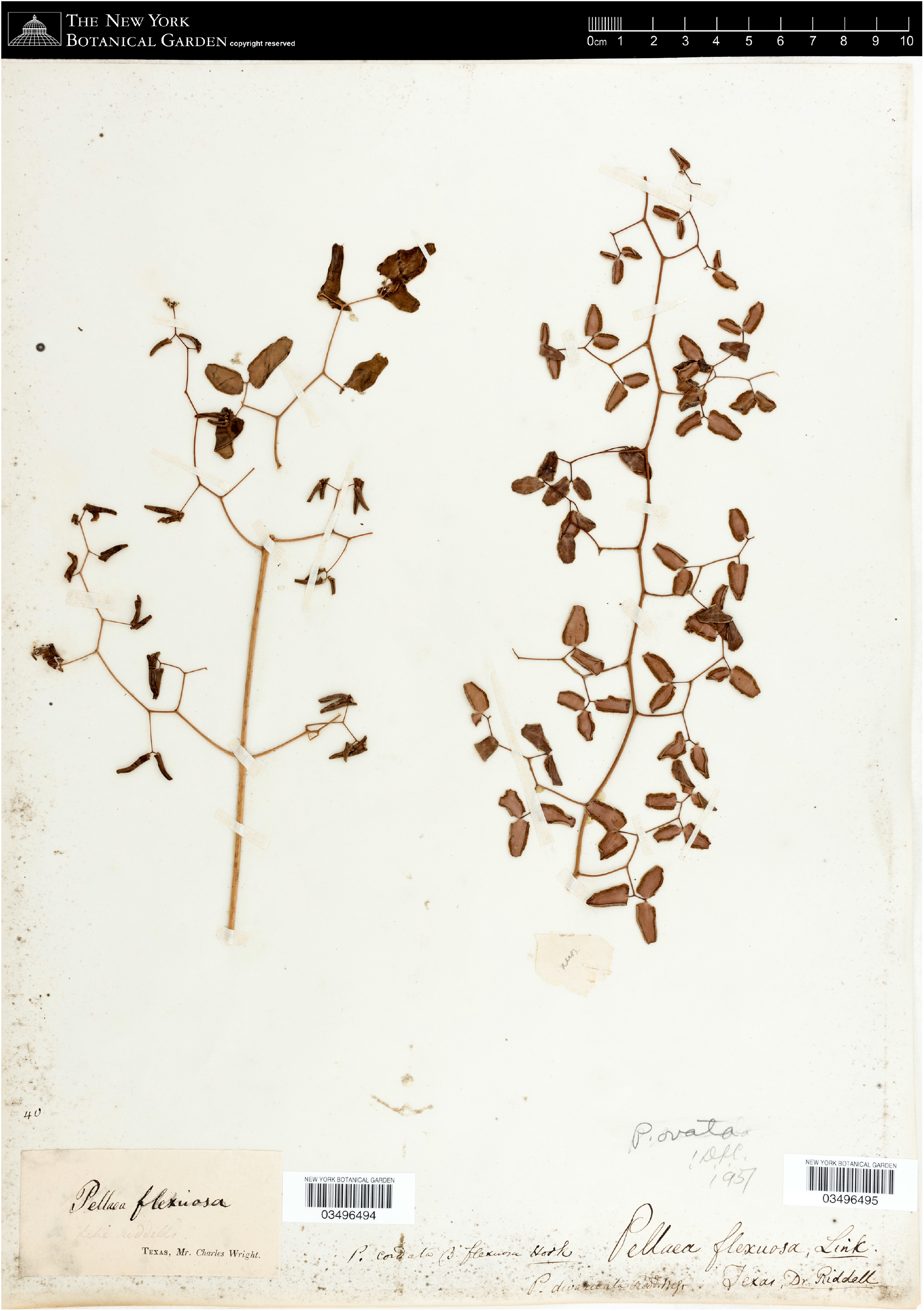
Type of *Pteris zygophylla*: *Riddell s*.*n*., Oct 1839, Comanche country, Texas (ny 3496495). This and other ny images belong to The C. V. Starr Virtual Herbarium (http://sciweb.nybg.org/VirtualHerbarium.asp).

**Figure 4.**
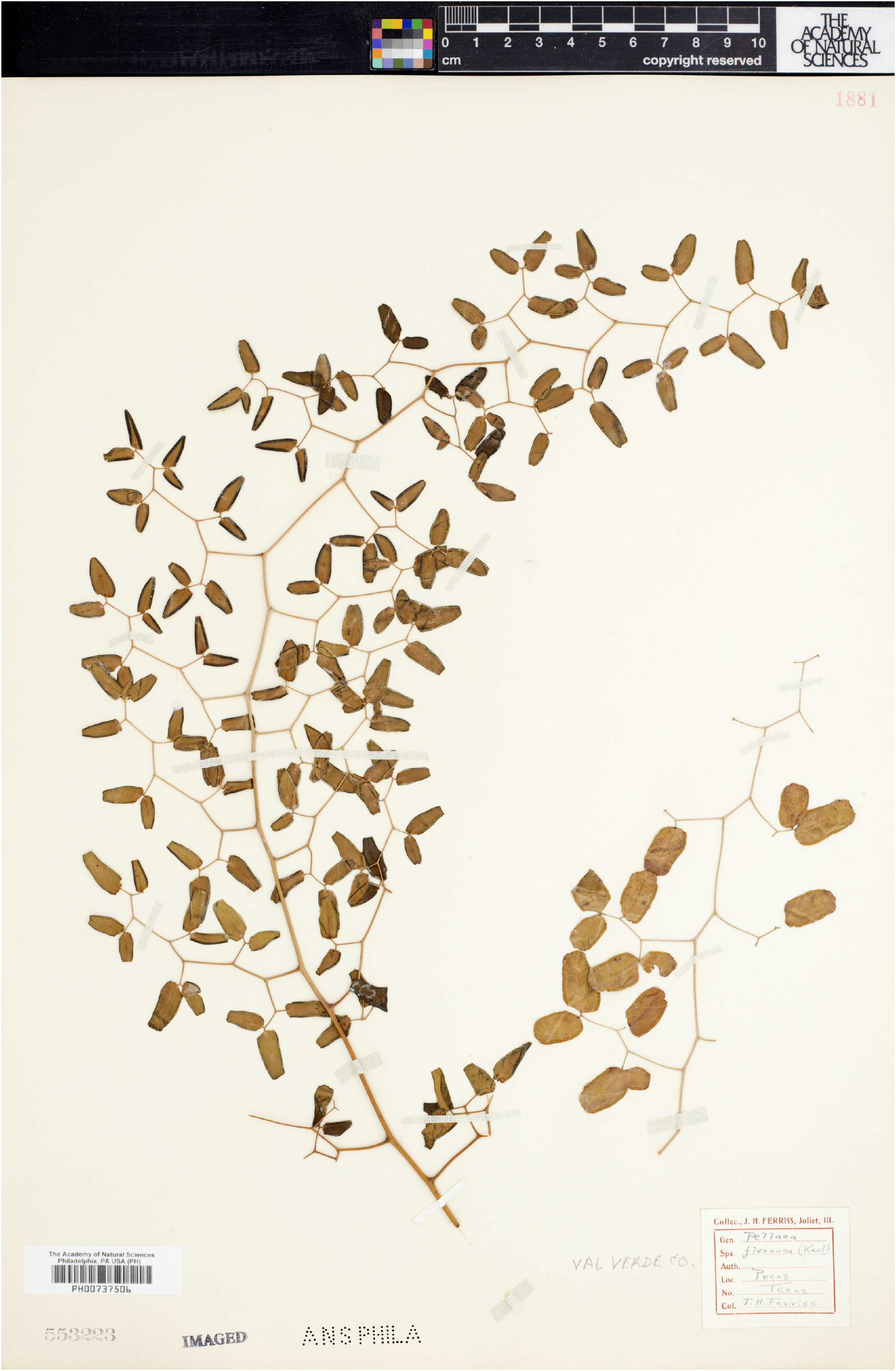
*Ferriss s*.*n*., s.d., Pecos [River], Val Verde Co., Texas (ph 737506).

**Figure 5.**
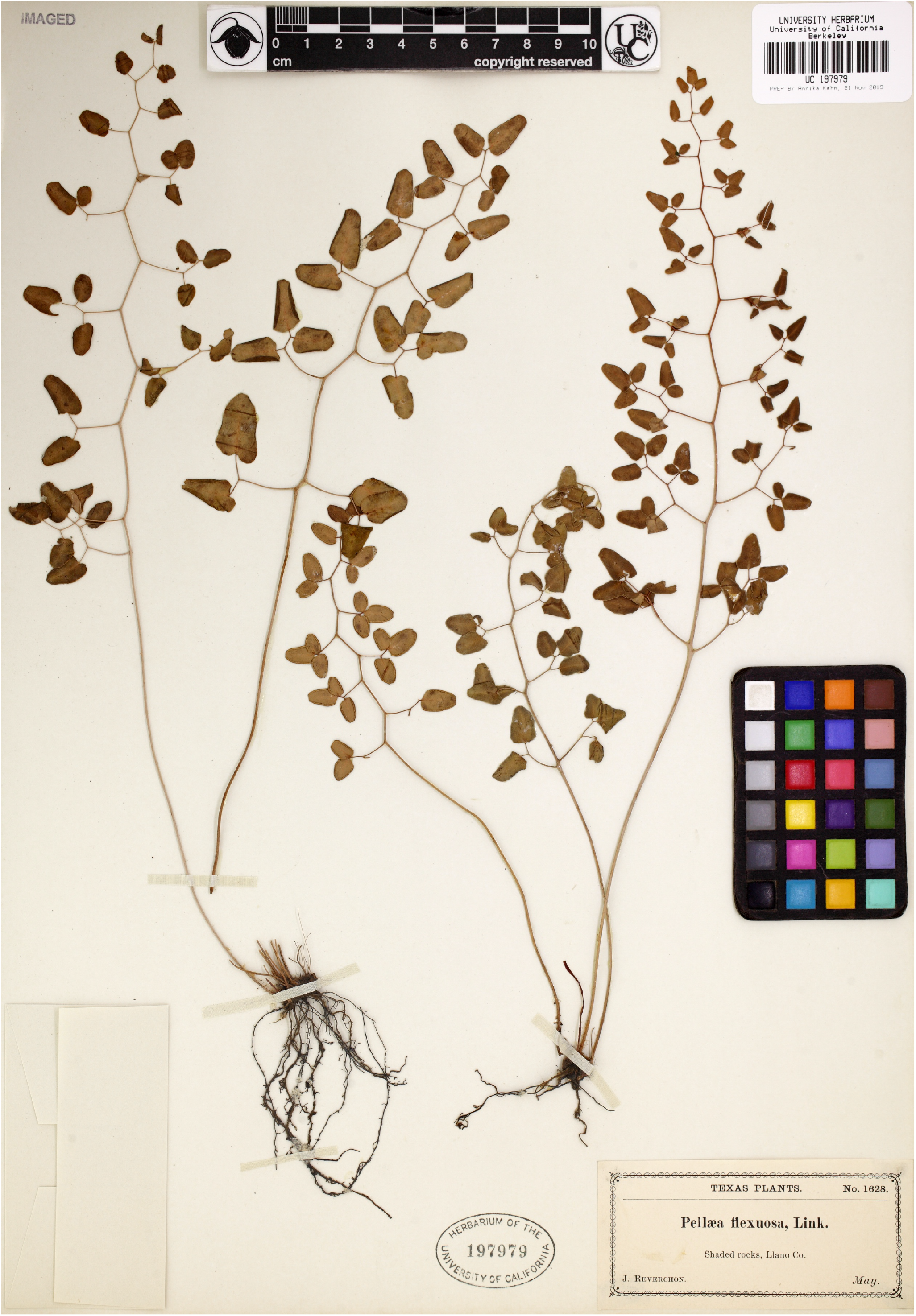
*Barbara s*.*n*., 6 Feb 1954, Inks State Park, Burnet Co., Texas (uc 2076829).

**Figure 6.**
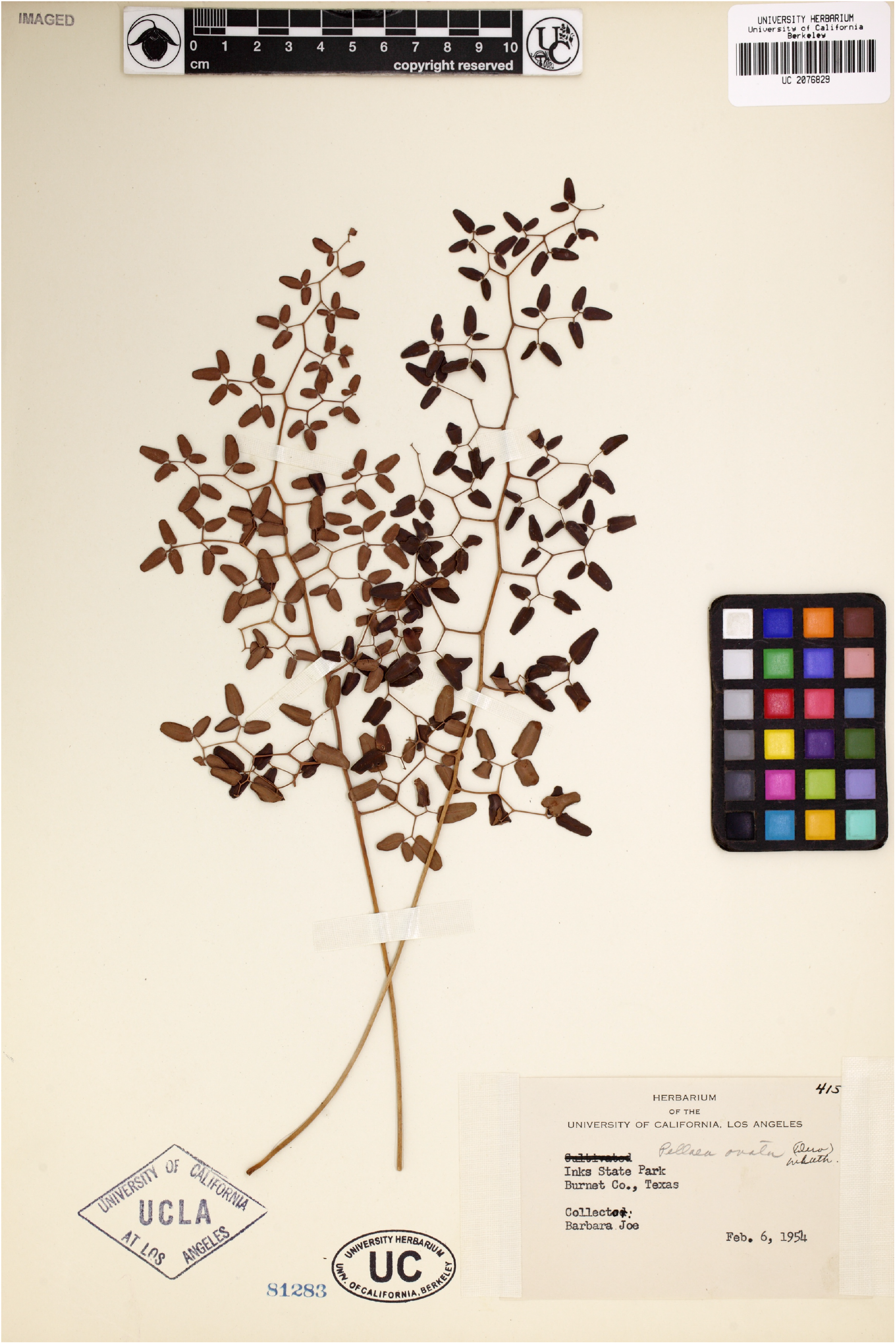
*Reverchon 1628*, May [1885], Llano Co., Texas (uc 197979).

**Figure 7.**
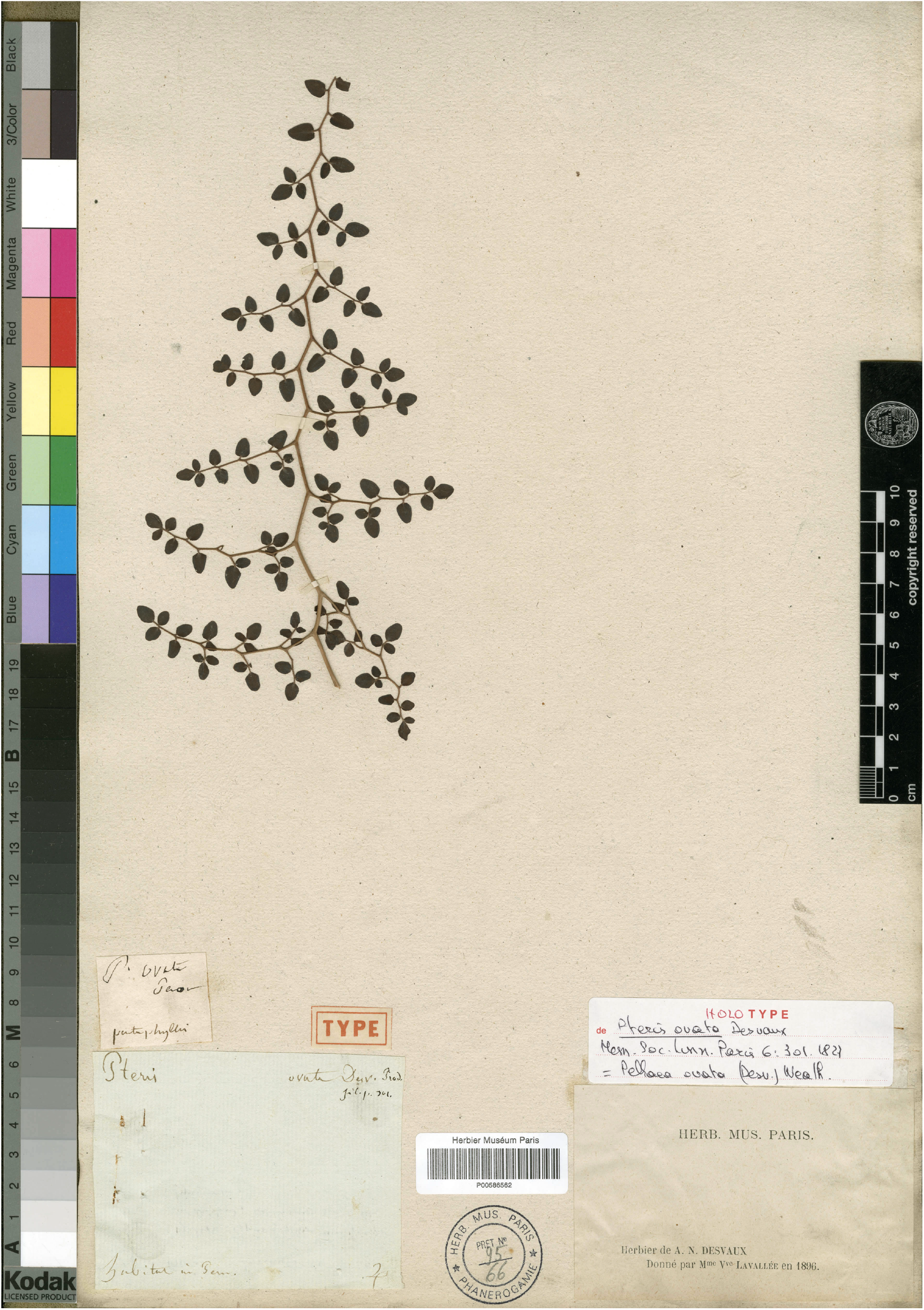
Type of *Pteris ovata*: *Anonymous, s*.*n*., s.d., Peru (p 586562).

Rhizomes creeping, slender, 2–3 mm in diameter; *scales* loosely appressed, lanceolate, 2–3 × 0.5–0.8 mm, bicolorous, centers lustrous black, with thin, brown, pectinate to erose-serrulate margins. Leaves 30–120(–200) × (7–)10–30(–40) cm, ascending to sprawling, often subscandent; *stipe* 0.5–0.9 times as long as the blade, rounded or flattened adaxially, scaly for 0.5–2 cm at the base, basalmost scales persistent, dense, like those of the rhizome, more distal scales gradually deciduous, sparse, pale, and linear; otherwise glabrous or becoming sparsely puberulent near the base of the blade; stipe and rachis tan to light reddish-brown, turning light gray with age; *rachis* weakly to strongly flexuous, often nearly straight toward the base of the blade and becoming strongly flexuous distally, puberulent distally or throughout; *blade* lanceolate, usually 3-pinnate, sometimes 2- or 4-pinnate, with (5–)8–15(–20) pairs of pinnae, alternate, occasionally subopposite and becoming alternate distally or (rarely) subopposite throughout, distalmost 5–7 pinnules borne singly on the rachis. Pinnae reflexed to slightly ascending, sometimes gently arcing toward the apex of the leaf, the larger pinnae usually 2-pinnate, with 9–50(–65) pinnules borne singly, central axis obvious and not appearing dichotomous, sometimes the ultimate two pinnules paired and very unequal in size; *costae* weakly to strongly flexuous or (rarely) straight, base reflexed to horizontal or occasionally weakly ascending, puberulent throughout or at least in distal half; *stalks* of the pinnules 1–5(–8) mm, densely puberulent, trichomes 0.1–0.3 mm long, dull tan to pale reddish-brown; costae &stalks of the pinnules the same color as the rachis but darkening distally. Pinnules ovate or (rarely), broadly lanceolate, 5–14(–21) × 3–10(–15) mm, 1.5–2(–2.5) times longer than wide, subcoriaceous, veins indistinct or occasionally distinct abaxially, glabrous or (rarely) sparsely pubescent on one or both surfaces; *base* cordate, truncate, or rounded, sometimes incised only immediately around the stalk of the pinnule, usually a little oblique on terminal pinnules but equilateral on lateral pinnules; *apex* rounded or rounded-acute, sori usually converging at the apex or with a relatively narrow and inconspicuous gap between them or (rarely) the apex rounded-truncate and with a wide gap between the sori; *sori* often visible adaxially as a swelling toward the pinnule margin, false indusia 0.3–0.6 mm wide, revolute, entire, little differentiated from the rest of the pinnule.

32 spores per sporangium; plants apogamous, triploid. Tryon &Britton (1958) and Tryon (1968) report apogamous triploids, n = 3n = 87, from Mexico (*Correll &Gentry 22792, Tryon &Tryon 5134*; identity verified from images). Tryon also counted spores and found only 32-spored plants from southern Mexico, Central &South America, and Hispaniola. A sexual diploid form may also exist. Velázquez-Montes (2018) reports the species 64-spored in Guerrero based on a specimen with abaxial pinnule surfaces sparsely pubescent with jointed trichomes (*Lorea 1445*, fcme; not seen). A sexual diploid count from Costa Rica is mentioned by Mickel &Smith (2004) and attributed to Gómez-Pignataro (1971; I have not seen the paper) in the Chromosome Counts Database (Rice &al. 2014).

Northwestern Mexico (Sonora &Baja California Sur), south to northern Argentina (Catamara), east to northern Venezuela (Caracas), and with a disjunct population in southern Brazil (São Paulo). Map 1 &Map2. Mostly subtropical highland climates, in seasonally dry woodlands on varied geology.

Argentina. Catamarca: *Castillon s*.*n*. (mich 1208290, u 1040827). Jujuy: *Cockerell s*.*n*. (us 1231030); *Eyerdam &Beetle 22417* (uc 652335). TucumÁn: *Schreiter 1515* (u 1040828); *Venturi 10367* (us 1694546); *Venturi 1246* (us 1694311).

Bolivia. Chuquisaca: *Kessler &al. 4915* (us 3366991). Cochabamba: *Cárdenas 3313* (f 660505). *Cárdenas 4798* (us 2135362); *Kessler &al. 9595* (uc 1620788); *Kuntze s*.*n*. (ny 3902561 &3902565). La Paz: *Brooke 5509* (f 660504, ny 3902583, u 1040812); *Feuerer 5794a* (f 660503); *Kessler &al. 10380* (uc 1621412); *Lewis 35136* (f 660502, ny 3902589, uc 1585093, us 3218330); *Lewis 35402* (uc 1585109 &1585110); *Rusby 142* (ny 3902568, us 1069655). Santa Cruz: *Nee 58640* (ny 3527925). Tarija: *Krapovickas &al. 19172* (uc 1383217).

Brazil. SÃo Paulo: *Prado 1658* (ny 2422515 &2422515).

Colombia. Cauca: *Anonymous B*.*T*.*772* (ny 3902563). Cundinamarca: *Haught 6053* (us 2016847 &2016848). NariÑo: *Garganta s*.*n*. (f 660473). Santander: *Killip &Smith 16382* (us 1352128); *Killip &Smith 17440* (us 1353040); *Killip &Smith 19090* (f 660472, us 1354391); *Killip 16382* (ny 3902560); *Killip 19090* (ny 3902566). Valle del Cauca: *Cuatrecasas 20467* (f 660474, us 2018934).

Costa Rica. Alajuela: *Brade 16403* (us 472486). Cartago: *Brade 199* (ny 3902419, uc 403629); *Standley &Valerio 49534* (us 1308311); *Valerio 196* (us 1316803). Heredia: *Gómez 2776* (f 633218).

Ecuador. Carchi: *van der Werff &Gudiño 10654* (uc 1583356). Chimborazo: *Camp E-3167* (uc 951273). Imbabura: *Baker 7356* (ny 01527756); *Mexia 7404* (uc 619500); *Mexia 7426* (uc 619486). Loja: *Fay 4505* (ny 3902581, uc 1744211). Pichincha: *Sodiro 3/908* (uc 1193456).

Guatemala. Chiquimula: *Steyermark 31418* (f 633206, us 1793000). Guatemala: *Dziekanowski &al. 3139* (wis 374440); *Dziekanowski &al. 3457* (wis 374439, mich 1208301). Huehuetenango: *Molina 21333* (f 633201); *Molina 30250* (f 633202); *Standley 81200* (f 633205, us 1840423); *Steyermark 48110* (f 633211, us 1917108); *Williams &al. 22029* (ny 3902416, us 2425614); *Williams &al. 22325* (us 2425498). Jalapa: *Standley 77095* (f 633208). SacatepÉquez: *Standley 58044* (f 633203); *Standley 80980* (us 1840418). SololÁ : *Hatch &Wilson 291* (brit 497255 &497256, us 1687952); *Hatch &Wilson 330* (brit 497253, uc 755792 &755792, us 1687974); *Hatch 293* (f 633200); *Steyermark 47124* (f 633207); *Steyermark 47282* (f 633210, us 1917068).

Honduras. Comayagua: *Standley 56496* (f 633214, us 1309261).

Mexico. Aguascalientes: *McVaugh &Koelz 123* (mexu 560200, mich 1508445). Baja California Sur: *León 3415* (uc 1577802). Chiapas: *Alava 1312* (mexu 157691, uc 1094408); *Alava 1342* (mexu 157692, uc 1094426); *Breedlove 39908* (mexu 246525); *Méndez 9180* (mexu 996146); *Najarro &Moreno 2324a* (mexu 1432485). C hihuahua : *Correll &Gentry 22792* (msc 267091, uc 1225019, us 2359092); *Gentry 1538* (uc 576781). C iudad de México: *Lyonnet 861* (us 1821451); *Rzedowski 24259* (mich 1208294, msc 267088). D urango : *Corral-Díaz &Worthington 67* (ind 3410); *McGill &al. 9406* (asu 3716, des 7379). Guerrero: *Hinton &al. 11305* (us 1792131); *Valencia 1084* (mexu 1004800). H idalgo : *Broun s*.*n*. (ph 737296); *Gastony &Yatskievych 86-42* (ind 3411); *Gimate 6* (huap 24456); *González 3249* (mich 1208306); *Hernández &Hernández 4573* (mexu 317554); *Matuda 32507* (mexu 762304); *Parfitt &al. R-6004* (asu 3717). J alisco : *Díaz 3265* (mexu 183122); *Díaz 8790* (uc 1478450); *Díaz 968* (uc 1440040 &1440040); *Jones s*.*n*. (rsa 32224); *Judziewicz &Guzmán 5071* (wis 374443); *Mones 18213* (uc 1534803); *Pringle 2032* (ny 3902367); *Pringle 5408* (us 961231, vt 194455); *Pringle s*.*n*. (uc 679354); *Pringle s*.*n*. (uc 150612, vt 194451); *Rose &Painter 7594* (ny 3902393, us 451204); *Santana 7268* (brit 432137, wis 374442); *Vázquez &al. 13013* (wis 374444). México: *Borgeau 251* (uc 1194183); *Dorantes-Hernández &al. 148* (mexu 1403825); *Goodding 2180* (uc 163465); *Ledesma 1823* (mexu 1380612); *Rzedowski 27948* (mich 1208233, wis 374449); *Tejero 2137* (mexu 1182955); *Tryon &Tryon 5134* (us 2425788). M ichoacÁn : *Arsène 3645* (us 1030130); *Arsène 5496* (us 1000012); *Arsène 6567* (us 1030126); *Arsène 9984* (us 1000010); *Arsène 9985* (us 1000013); *Contreras 77* (mexu 1452407); *Cowan &al. 5677* (wis 374450); *Manuel 1662* (mexu 676773); *Pérez &al. 2189* (mexu 874538); *Nelson 6546* (us 399138); *Salazar &al. 9200* (mexu 1399537 &1399538); *Tejero &Sánchez 4755* (mexu 1306828); *Yatskievych 86-31* (ind 3407). Morelos: *Lyonnet 521200002* (mexu 648281 &648282). Nuevo LeÓn: *Dorr &al. 2575* (mexu 357554 &365429, uc 1504397); *Mueller &Mueller 1130* (mich 1208221); *Pennell 17229* (ph 737294). Nayarit: *Tellez 12839* (mexu 547652). Oaxaca: *Camp 2235* (ny 3902408); *Camp 2487* (ny 3902409, uc 1507643); *Conzatti &al. 3030* (us 794648); *Figueroa &Guzmán 516* (mexu 1340865); *Galeotti 6558* (yu 20019); *Gastony &Yatskievych* 86-37 (ind 3408); *Gastony 86-38* (ind 3414); *Gereau &Saynes 2137* (mo 12396); *Ibarra &al. 133* (mexu 1363073); *Knobloch 2204* (msc 267093); *Mendoza &al. 429* (mexu 1353749); *Mickel &Hellwig 3706* (ny 3902407, uc 1494872); *Mickel &Hellwig 3847* (ny 3902362, uc 1493838); *Mickel &Hellwig 3898* (ny 3902410, uc 1494873); *Mickel &Leonard 4508* (ny 3902411, uc 1503049); *Mickel &Leonard 4959* (ny 3902400, uc 1466768); *Mickel &Pardue 6475* (ny 3902412, uc 1466769); *Mickel 3922* (ny 3902414); *Mickel 4488* (ny 3902413); *Mickel 6650* (ny 3902406); *Mickel 754* (mich 1208305); *Mickel 830* (us 2420358); *Mickel 886* (us 2420303); *Smith 2057* (ny 3902388, us 312920); *Sundue s*.*n*. (vt 194443); *Yatskievych &González 85-210* (ind 3413); *Yatskievych &Gastony 89-282* (ind 3415). Puebla: *Arsène 1477* (us 1030122); *Arsène 1620* (us 1030129); *Arsène 297* (us 1030124); *Arsène 539* (us 1030125); *Arsène 9955* (us 1030127); *Arsène 9956* (us 1030128); *Arsène 9957* (us 1030123); *Arsène s*.*n*. (mich 1208220); *Arsène s*.*n*. (ncu 432685, ph 737288, uc 2017349); *Cerón &Coombes 6738* (huap 60680); *Cerón 1543* (huap 56697); *Cerón 2371* (huap 65888); *Cerón 2498* (huap 67347); *Cerón 2635* (huap 67414); *Cerón 378* (huap 27834, mexu 1403971); *Copeland 108* (mich 1208228 &mich 1208304, msc 267087, ny 3902361, uc 600928); *González 7701* (huap 64072); *Purpus 1152* (uc 150539); *Purpus 4035* (uc 150302); *Sanchez-Ken 306* (mexu 520721). San Luis PotosÍ: *Pringle s*.*n*. (mich 1208292, us 2258280, vt 194444). Sonora: *Ferguson 2962* (mo 3129089); *Reina &Van Devender* 97-448 (mexu 898129). a: *Calzada 4266* (uc 1533410); *Hernández &Chacón 472* (uc 1543512); *Lemmon &Lemmon 333* (uc 156714); *Matuda 1173* (mexu 88787, mich 1208296); *Matuda 198* (mich 1208303); *Mohr 48* (yu 20028); *Seaton 39* (ny 3902364).

Nicaragua. Jinotega: *Standley 10192* (f 633217); *Standley 9792* (f 633216); *Standley 9821* (f 633215); *Stevens &Montiel 29530* (mo 100256001).

Peru. Amazonas: *Hutchinson &Wright 4891* (uc 1200111); *van der Werff &al. 14659* (uc 1728858). ApurÍmac: *Anonymous s*.*n*. (uc 565290); *Nuñez 7194* (ny 3902571); *Stork &Horton 10712* (uc 656918); *Vargas 8774* (uc 935592). Cajamarca: *Dillon 4536* (f 660480, ny 3902586); *Sagástegui 14689* (ny 3902588); *Sagástegui 14816* (ny 3902587). ContumazÁ : *Sagástegui &al. 15892* (uc 1732111). Cusco: *Galiano 5510* (uc 1870662); *Suclli 2175* (uc 1978767); *Valenzuela 4586* (uc 1978480). Huancavelica: *Hutchinson 1685* (uc 1210536). Huanuco: *Woytkowski 34255* (uc 1015524). JunÍn: *Coronado 243* (uc 1052646). La Libertad: *Bussmann &al. 16849* (mo 100386676); *Bussmann &al. 17346* (mo 100546970); *Bussmann &al. 17410* (mo 100547272); *Bussmann &al. 18475* (mo 100666202).

Venezuela. Aragua: *Fendler 89* (yu 20063). Caracas: *Vogl s*.*n*. (uc 404685). FalcÓn: *van der Werff &Wingfield 7453* (huap 27834, mexu 1403971). MÉrida: *Ortega &Díaz s*.*n*. (huap 27834, mexu 1403971). TÁchira: *Ortega &van der Werff 2878* (uc 1524786, ny 3902569 &3902573).

Typification of *Pteris flexuosa*—The protologue from Schlechtendal &Chamisso (1830) follows:

> 785 Pteris *flexuosa* Kaulf. mspt. in hort. berol. Rachide insignius flexuosa magisque pubescente vix satis a *Pteride cordata* Sw. diversa. W. spl. pl. p. 392, herb. no. 20005. (spec. Humb.), HBK 1. p. 15, a qua non differt *Pt. sagittata* W. herb. no. 20006. (spec. Humb.) HBK 1. p. 14.—In sylvis prope Jalapam. Aug.

There are seven sheets in online databases to consider; none match the protologue fully. No question of taxonomy hinges upon the identification of the type, as all are identifiably *Pellaea ovata* s.s. They bear Schiede’s name alone, despite attribution to Schiede &Deppe by Schechtendal &Chamisso. All seven are original material as defined by ICNafp Art. 9.4. Neither the protologue nor sheet labels allow any single sheet to be identified as the holotype. I recognize as types a set of four sheets united with the protologue by annotation, with each other by a mix of annotation &label information. Two, hal 137767 &le 8610, are annotated “785 *Pteris flexuosa* Kaulf.” in Chamisso’s hand. I identify Chamisso from hal 81851, in the same set of Schiede’s specimens &referenced in the same work. It bears a label “787 *Pteris pulchra* n. sp.” in a hand matching hal 137767 &le 8610, and is annotated “scripsit: A. v. Chamisso”. A third sheet, b 20 0103147, bears “785 *Pteris flexuosa* Kaulf. mspt.” in another hand. Following Heuchert &al. (2017), I take “785” to be an enumeration by Schlechtendal &Chamisso rather than Schiede’s collection number, and refer to these sheets as *Schiede s*.*n. “785”*. Although “785” would link them more strongly to each other were it Schiede’s, as Schlechtendal &Chamisso’s number it provides an unambiguous link to the protologue. A fourth sheet, hal 137766, has an original label with identical information to hal 137767. None of these four match the protologue’s “in sylvis prope Jalapam, Aug.” Instead, those with complete labels are marked “in dumetis Jalapae, May”. I take this to be an error that does not supersede authorial intent established by annotation &enumeration.

Of the seven potential types, only *Schiede 731* (hal 137765) has “in sylvis prope Jalapam, Aug.” Based on Heuchert et al. (2017), the number “731” does not conflict with the protologue. However, the sheet did not bear the name “*Pteris flexuosa*” until a recent, printed label. Although this could reasonably be considered a syntype, I cannot follow Heuchert &al. in identifying this sheet as a type to the exclusion of those to which the name was directly applied by the authors. The remaining two potential types (b 20 0103148 &hal 133764) say only “Mexico, Schiede” &not “*Pteris flexuosa*”. They must be either *Schiede s*.*n. “785”* or *Schiede 731*. Without grounds to assign them to one collection or the other, they are syntypes if hal 137765 is and in limbo otherwise.

Two sheets (b 20 0103147 &b 20 0103148) were annotated as holotypes by Palacios-Ríos in 1996. With seven sheets of original material to choose from and these two with a weaker claim than others (neither annotated by Chamisso nor matching the protologue), I do not see how either could be the sole specimen indicated or used.

## Discussion

Even in immature or fragmentary material *Pellaea ovata* s.s. and *Pellaea zygophylla* are reliably distinct. In good material they can be distinguished at a glance. Morphological ambiguity within *Pellaea ovata* s.l. revolves, instead, around *Pellaea oaxacana*. I think it is more likely that *Pellaea ovata* s.s. and *Pellaea oaxacana* are conspecific than that either is conspecific with *Pellaea zygophylla*.

The flexuous, pubescent rachides &costae of *Pellaea ovata* are usually sufficient to distinguish it from *Pellaea oaxacana*. However, while previous treatments uniformly describe *Pellaea ovata* as having flexuous rachides &reflexed pinnae, these characters are variable, always more pronounced distally. Also, descriptions of the pinnae are more accurately phrased as descriptions of the bases of the costae. The pinnae as a whole are more often perpendicular to the rachis or ascending than reflexed, especially in South American plants. The bases of the costae, though, are almost always reflexed in at least distal pinnae—usually conspicuously so, sometimes weakly. At the extreme, it may be only the distal bases of the costae that are slightly reflexed, and only the distal rachis and costae of distal pinnae that are weakly flexuous. These characters overlap with *Pellaea oaxacana*. I think pubescence distinguishes the two more consistently, but not all specimens can be confidently identified. Different botanists might easily draw the line between *Pellaea ovata* and *Pellaea oaxacana* in different places, depending on which features are emphasized.

*Pellaea ovata* is occasionally confused with *Pellaea intermedia* Mett. ex Kuhn. If relying on flexuosity of the rachis &reflexion of pinnae to distinguish them, this confusion should be rare but not entirely avoided. In uncertain cases, look for mature leaves of *Pellaea intermedia* to have rachides &costae bicolorous, pale &glabrescent adaxially, darker and conspicuously puberulent on the sides &abaxially. Although this is less evident on immature leaves and is not mentioned in published treatments of the genus, I believe it is a reliable marker for *Pellaea intermedia*. Russ Kleinman highlights it in his account of the species (on https://gilaflora.com as of Mar 2021).

The larger plants of this species are not easily accommodated by herbarium sheets, so the upper limits of leaf size given above are speculative. *Pellaea ovata* can become quite large.

> *Pellaea oaxacana* Mickel &Beitel
>
> Mem. New York Bot. Gard. 46: 271 (1988). Type: *Mickel 6279*, 11 Aug 1971, S of Sola de Vega, Oaxaca, Mexico (ny 144428!).
>
> Figure 1 (e, f), Figures 14–15.

**Figure 8.**
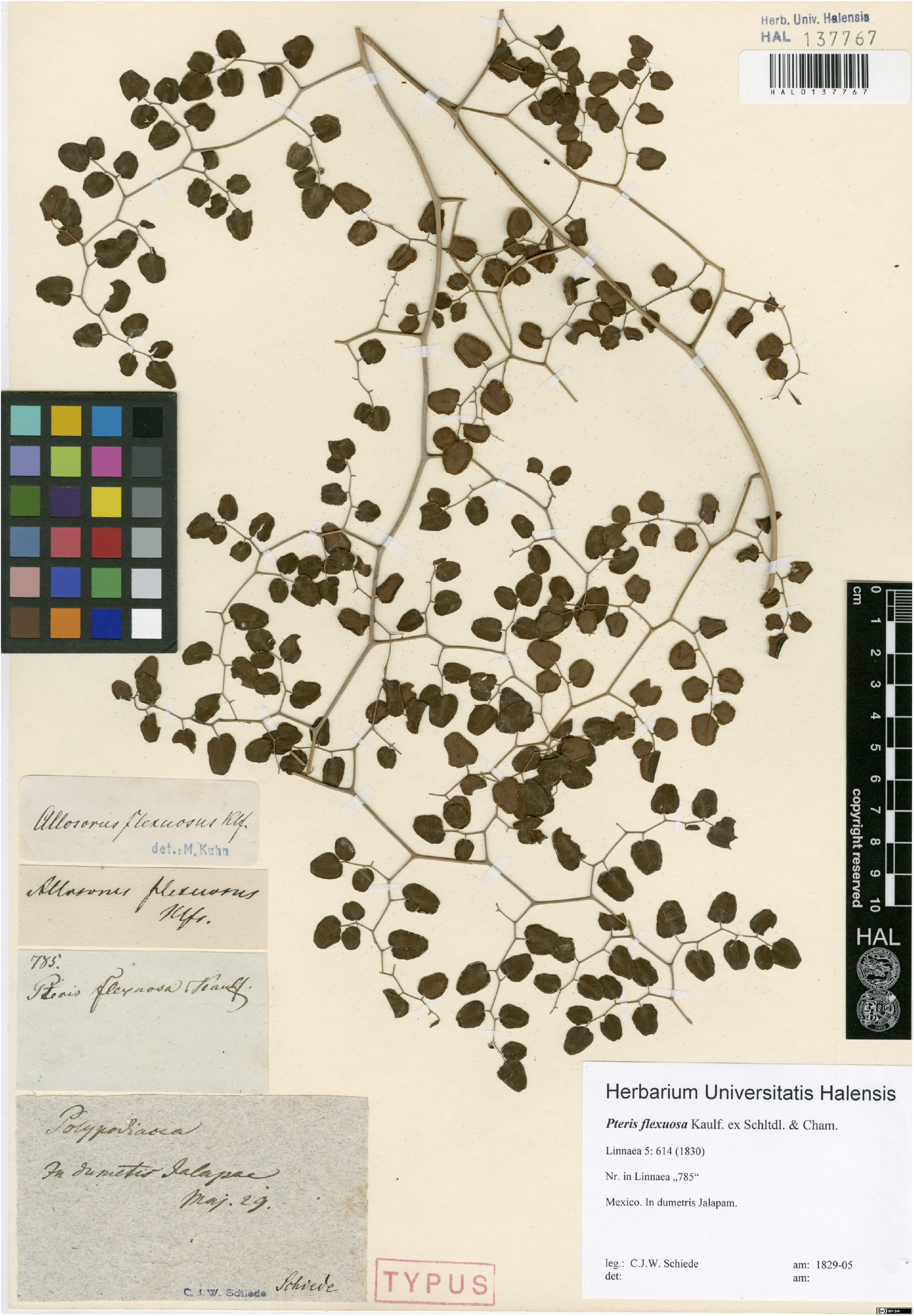
Type of *Pteris flexuosa*: *Schiede s*.*n. “785”*, May 1839, Jalapa, Mexico (hal 137767).

**Figure 9.**
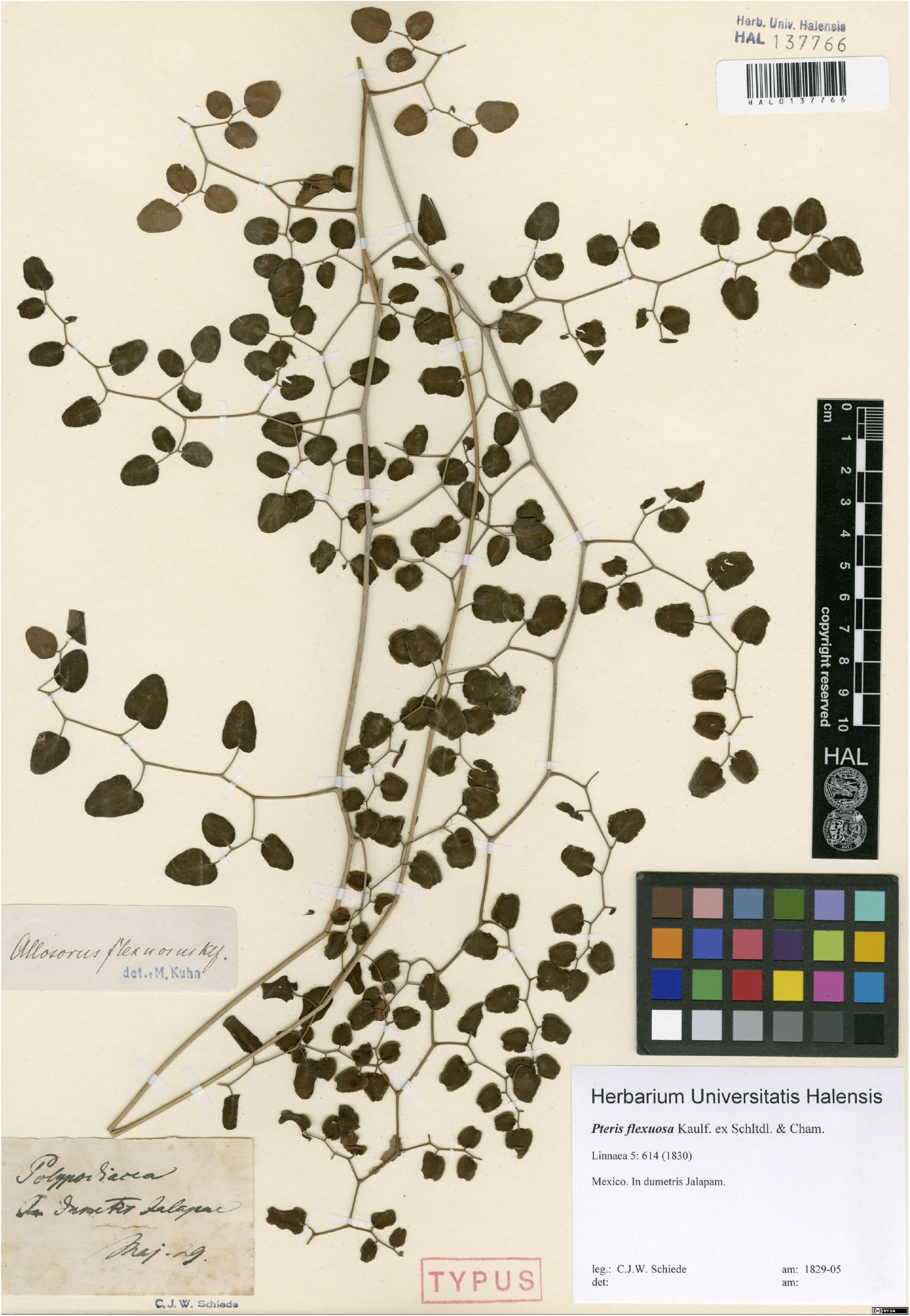
Type of *Pteris flexuosa*: *Schiede s*.*n. “785”*, May 1839, Jalapa, Mexico (hal 137766).

**Figure 10.**
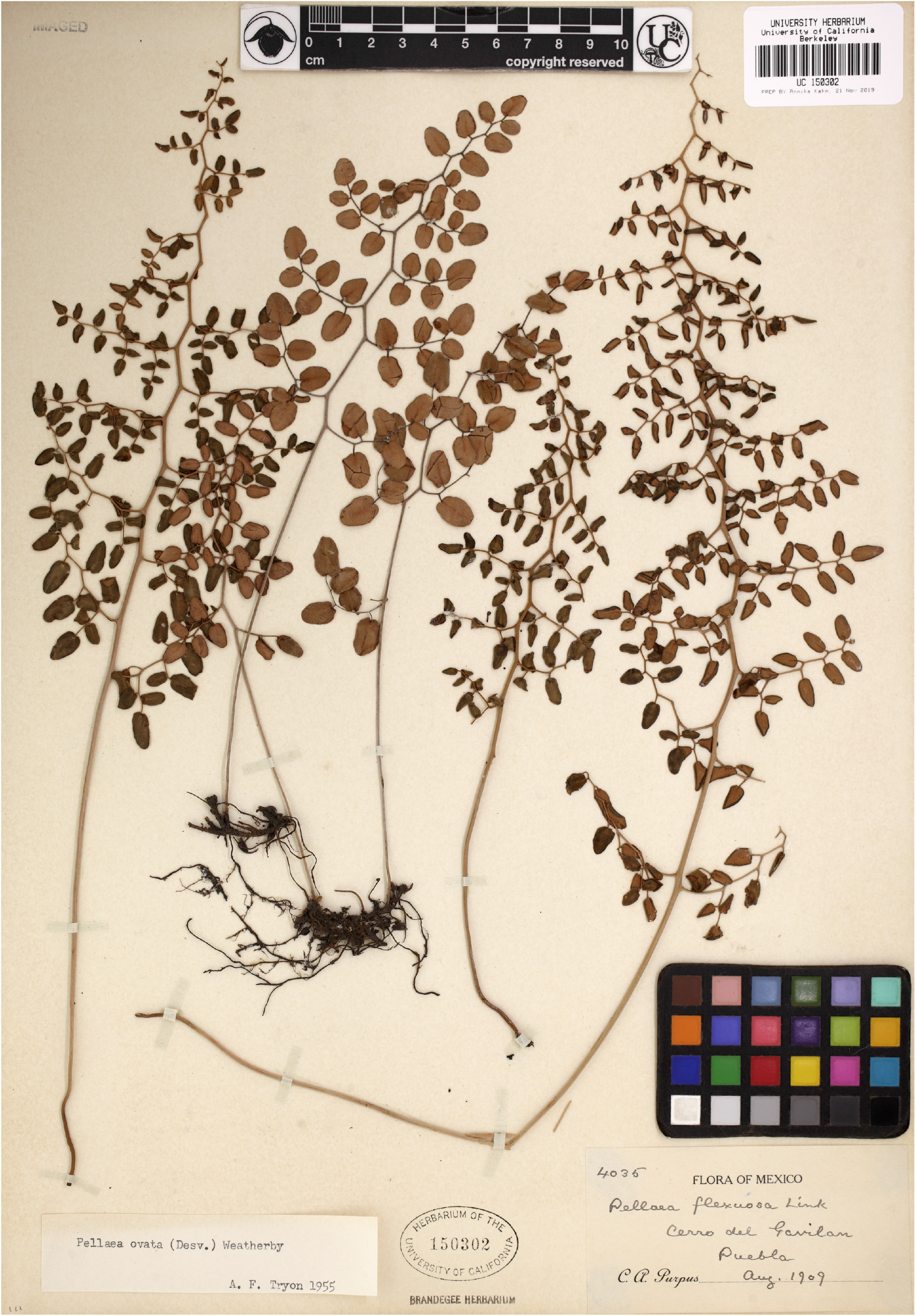
Type of *Pteris flexuosa*: *Schiede s*.*n. “785”*, May 1839, Jalapa, Mexico (b 20 0103148).

**Figure 11.**
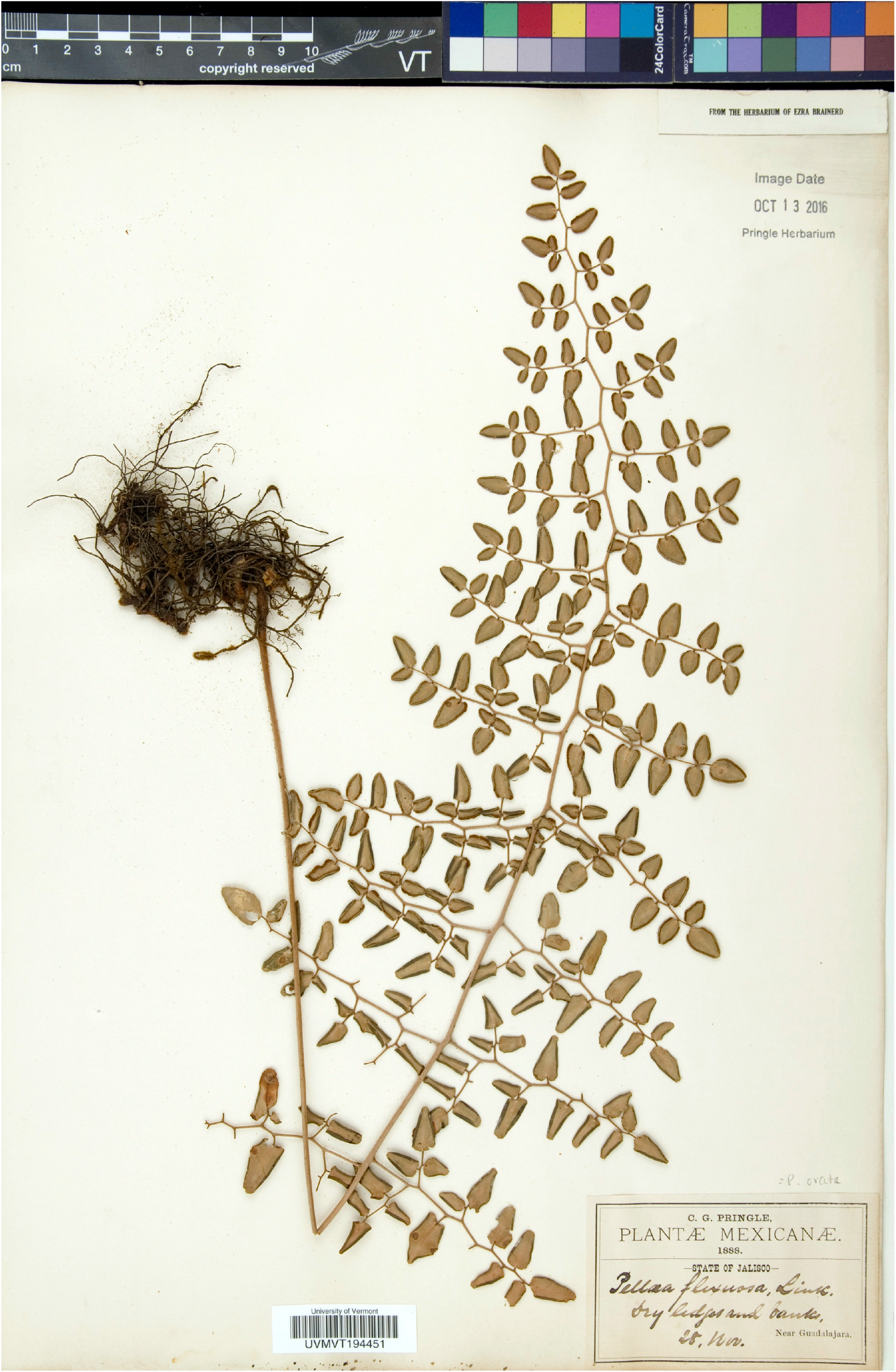
*Purpus 4035*, Aug 1909, Cerro de Gavilán, Puebla, Mexico (uc 150302).

**Figure 12.**
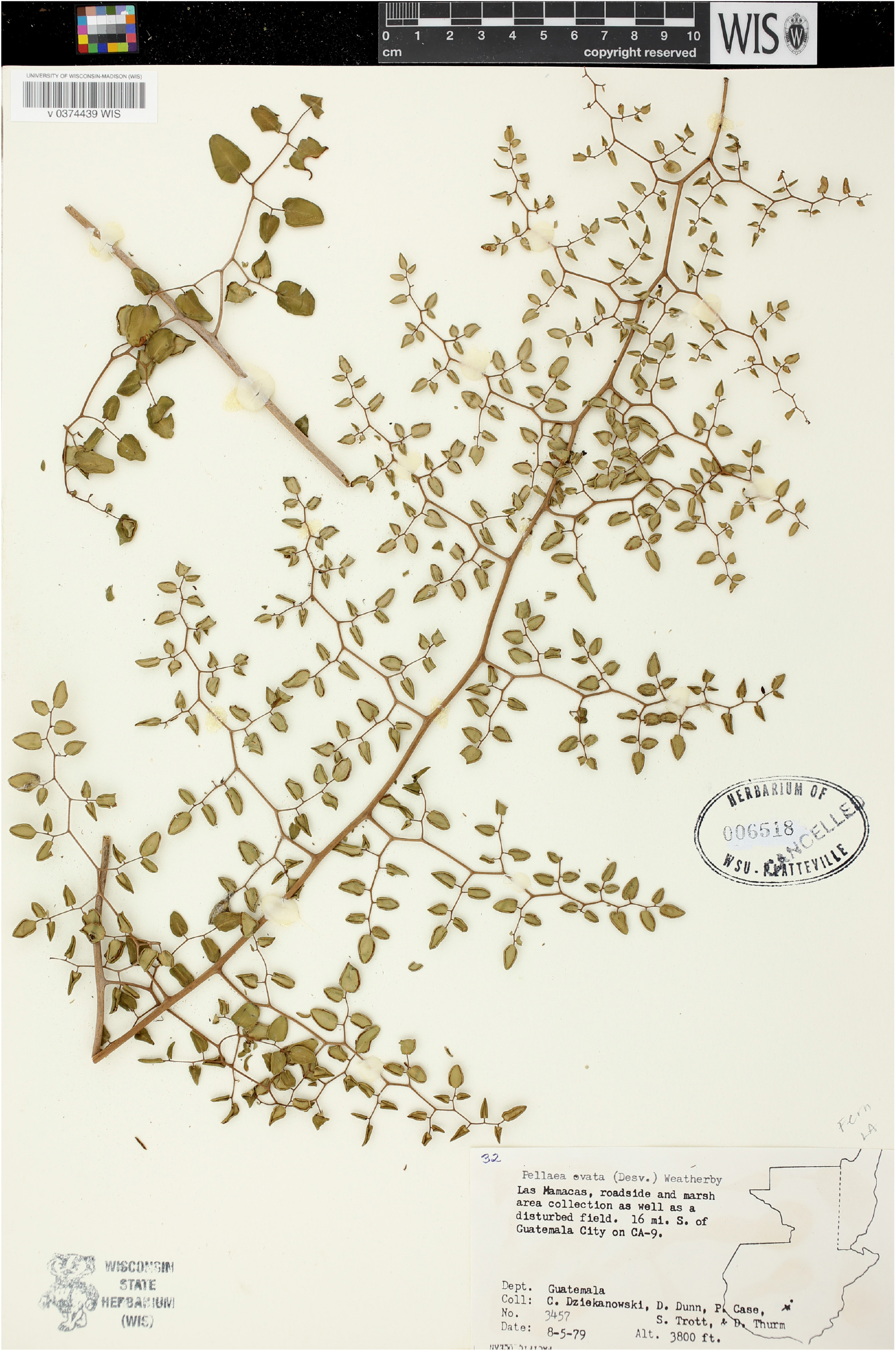
*Pringle s*.*n*., 28 Nov 1888, near Guadalajara, Jalisco, Mexico (vt 194451).

**Figure 13.** *Dziekanowski &al. 3457*, 5 Aug 1979, Las Mamacas, Guatemala (wis 374439).

**Figure 14.**
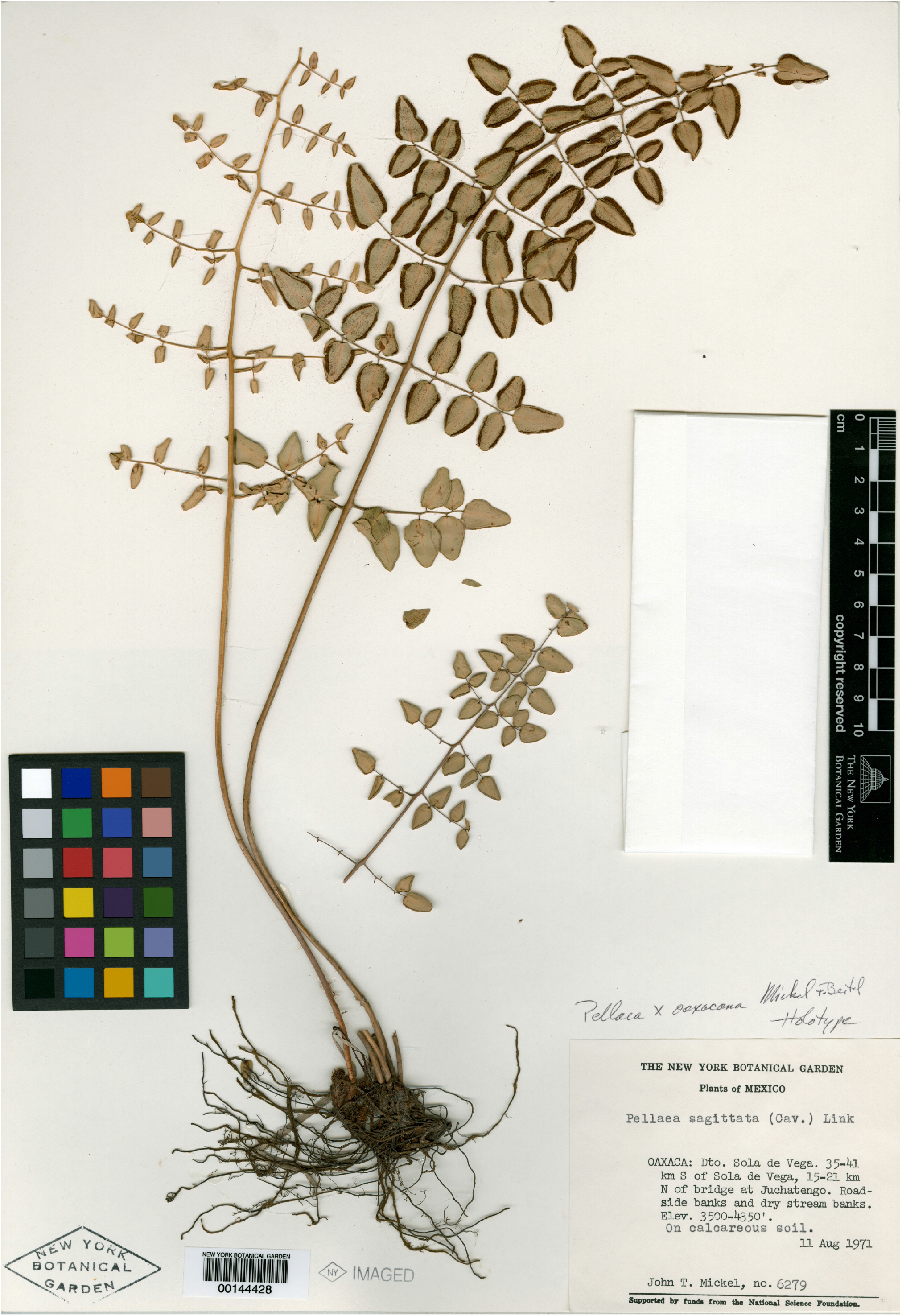
Type of *Pellaea oaxacana*: *Mickel 6279*, 11 Aug 1971, S of Sola de Vega, Oaxaca, Mexico (ny 144428).

Rhizomes creeping, slender, 1.5–3 mm in diameter; *scales* loosely appressed, lanceolate, 2–3 × 0.5–0.8 mm, bicolorous, centers dark reddish brown to dull black with wide, light brown, erose-denticulate margins. Leaves 20–60(–100) × 5–15(–30) cm, ascending to spreading, not subscandent; *stipe* 0.8–1 times as long as the blade, rounded or flattened adaxially, scaly for 1–3 cm at the base, basalmost scales persistent, dense, like those of the rhizome, more distal scales gradually deciduous, sparse, pale, and linear; stipe and rachis stramineous or tan, occasionally light reddish-brown, turning very pale gray with age; *rachis* straight throughout or becoming weakly flexuous distally, glabrous; *blade* lanceolate, usually 2-pinnate, 3-pinnate in large leaves, with 4–10 pairs of pinnae, usually subopposite, sometimes alternate or becoming alternate distally; distalmost 5–7 pinnules borne singly on the rachis. Pinnae stiffly horizontal or gently arced toward the apex of the leaf, pinnate, with 3–9 pinnules borne singly or, on exceptionally large leaves, 2-pinnate with up to 40 pinnules, branching in the pinnae never appearing dichotomous, *costae* usually not flexuous, occasionally weakly flexuous, especially in larger leaves, horizontal or weakly reflexed at the base, glabrous or occasionally sparsely puberulent distally; *stalks* of the pinnules short, 1–3(–6) mm, usually with a few translucent multicellular trichomes 0.1–0.3 mm long near the bases of the pinnules, sometimes sparsely puberulent throughout; costae &stalks the same color as the rachis but darkening distally. Pinnules rounded-trapeziform, ovate, or broadly lanceolate, (6–)8–20(–30) × (4–)6–12(–20) mm, 1.5–2 times longer than wide, coriaceous, glabrous, veins indistinct; *base* truncate to shallowly cordate, equilateral or slightly oblique on lateral pinnules; *apex* truncate or rounded, usually with a pronounced gap between the sori, at least on lateral pinnules, terminal pinnules more gently tapered, apices truncate to rounded-acute, sori sometimes converging at the apex; *sori* not visible adaxially, false indusia 0.3–0.7 mm wide, revolute, entire, thinning and becoming pale at the margin but otherwise little differentiated from the rest of the pinnule.

Likely 32 spores per sporangium and plants apogamous, triploid. Velázquez-Montes (2018) reports Guerrero plants to be 32-spored (based on *Carbajal 14*, fcme; not seen) but this appears to be the entire published record on the matter. Tryon (1968) reports plants from near San Luis Potosí with “leaf morphology closely resembling the sexual diploid type” (*Rollins &Tryon 58222*; not seen) to be apogamous, n = 3n = 87. *Pellaea oaxacana* has been found nearly this far north, the count may be attributable to it.

Southern Mexico and Guatemala, within a quadrilateral bounded by southern Nayarit, northern Hidalgo, southern Oaxaca, and southeastern Guatemala. Map 1 &Map 2. Subtropical highlands, mostly in seasonally dry woodlands, on limestone or igneous rocks.

Guatemala. Jalapa: *Standley 77618* (f 633204).

Mexico. Guerrero: *de la Rosa 583* mexu 1004799); *Nuñez 9697* (mexu 996106). Hidalgo: *Frye &Frye 2541* (uc 812396). Jalisco: *Barkley &al. 7618* (mexu 175312); *Barkley &al. 7647* (mexu 203022); *Garcia 55* (msc 267089); *Lemmon &Lemmon 145* (uc 111608); *Marker &Mellowes 108* (wis 374458); *Palmer 731* (yu 20018); *Santana &Sanchez 7052* (wis 374441). MÉxico: *Tejero 2606* (mexu 1182958, ny 3902334). MichoacÁn: *Vilas 20* (wis 374455). Morelos: *Rose &al. 10193* (us 453693). Nayarit: *Tellez 9330* (mexu 442736). Oaxaca: *Anonymous 40* (f 633263); *Anonymous s*.*n*. (f 633264); *Conzatti &González s*.*n*. (f 633135, central leaf, the others are *Pellaea cordifolia*); *Cruz-Espinosa 2003* (mexu 1052326 &1052327); *Hernández &Domínguez 105* (ny 1073135); *López 107* (mexu 1435515); *Mickel &Hellwig 3846* (mexu 859987, ny 3902339 &3902341, uc 1496055 &1728051, us 3124226); *Mickel &Leonard 4958* (uc 1493769 &1503046); *Mickel &Leonard 5001* (uc 1493770); *Mickel 4947* (ny 3902340); *Mickel 4958* (ny 3902337); *Mickel 5001* (ny 3902342); *Mickel 6251* (ny 3902338); *Mickel 6251* (uc 1494871); *Mickel 6279* (ny 144428); *Mickel 774* (mich 1208288, ny 3902343); *Rivera &al. 22* (mexu 1360553); *Salas &Sánchez 5015* (ny 3902335); *Santiago 16* (ny 3902333 &3902336); *Solheim &Powers 813* (wis1 374448). Puebla: *Purpus 4034* (uc 150522). Querétaro: *Arsène &Agniel 10649* (us 1032549); *Beck &al. 1241* (mo 3624678); *Rose &Rose 11195* (us 453977).

## Discussion

*Pellaea oaxacana* is quite distinctive in its “pure” form, with 2-pinnate leaves, rachides &costae glabrous and straight. Its distinction from *Pellaea ovata* is not always clear. This is especially true of larger plants of *Pellaea oaxacana*, which can be 3-pinnate and are particularly likely to have flexuous rachides &costae reflexed at the base. Only young plants with pinnate or barely 2-pinnate leaves are difficult to distinguish from *Pellaea zygophylla*.

*Pellaea oaxacana* can be confused with *Pellaea intermedia*, a mistake I made with plants in Gastony’s greenhouse whose labels had been lost or broken. Geography should be sufficient to distinguish them when the origin of the plant is known. Any lingering uncertainty should be resolved by the bicolorous rachides &costae of *Pellaea intermedia. Pellaea oaxacana* is also sometimes confused with *Pellaea sagittata* (see discussion of that species below). Except perhaps in very battered or fragmentary material, this can be resolved by looking for at least a few persistent scales throughout the length of the stipe and on the rachides &costae in *Pellaea sagittata*, while any scales above the basal several centimeters of the stipe in *Pellaea oaxacana* are quickly deciduous, gone well before a leaf is unfurled.

*Pellaea oaxacana* has not been reported previously from Guatemala, but *Standley 77618* matches the species in all respects. *Standley 77095*, collected nearby, has the rachides only slightly flexuous distally on some leaves, entirely straight on others; costae slightly reflexed at base, very slightly flexuous, and arced a little toward the leaf apex; but rachides densely puberulent distally; costae densely puberulent throughout, or glabrescent in the basal 1 cm on some of the larger &more basal pinnae. This pair nicely illustrates some of the difficulty in separating *Pellaea ovata* &*Pellaea oaxacana*. Since I place greater emphasis on pubescence, I identify *Standley 77095* as *Pellaea ovata*.

### *Pellaea ovata* s.l. on Hispaniola

Plants on Hispaniola do not fit comfortably in either *Pellaea ovata* s.s. or *Pellaea zygophylla*. They are excluded in the descriptions above and in the maps. Typical specimens are shown in Figures 16 &17. They are distinctive in the following combination of features:

**Figure 15.**
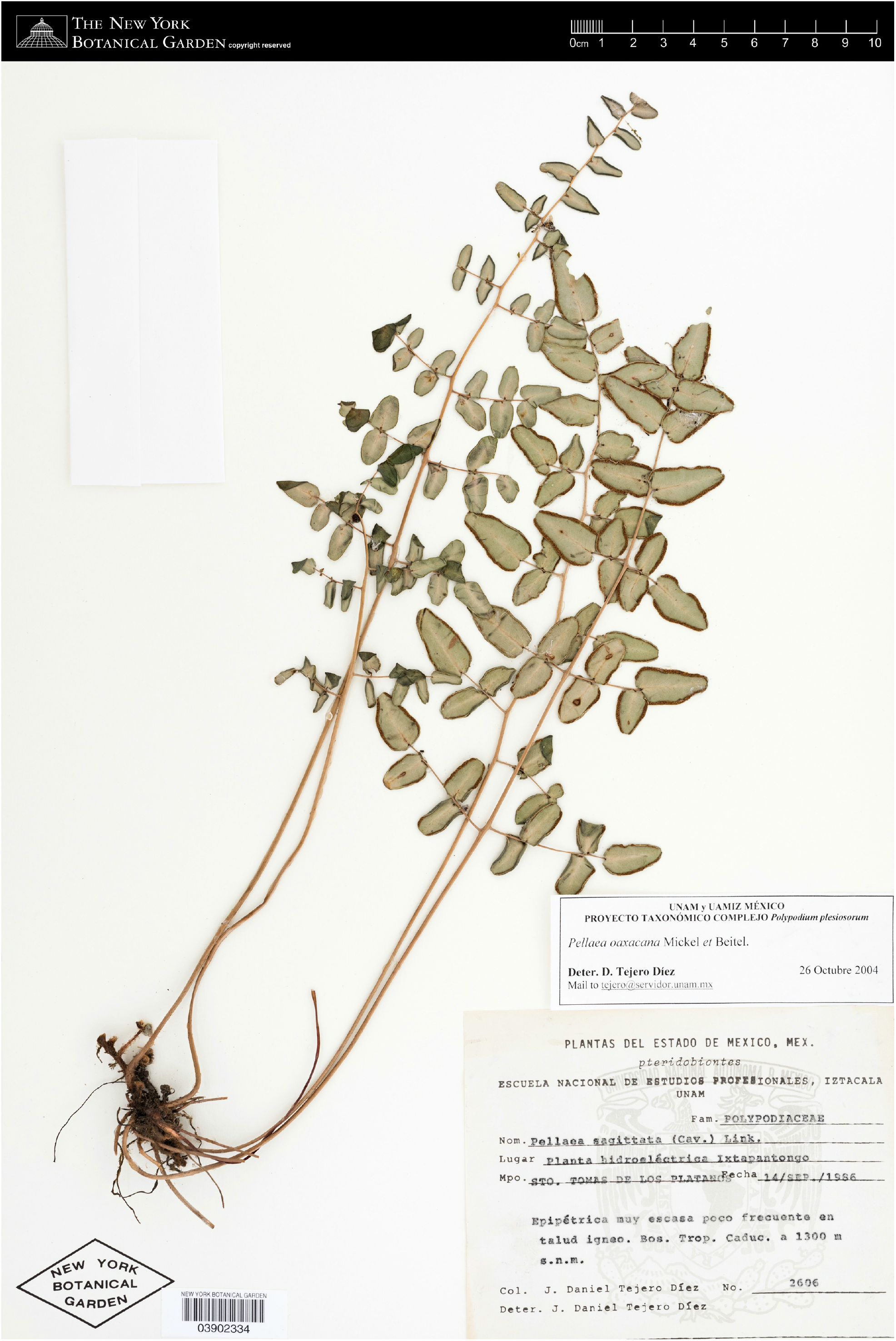
*Tejero 2606*, 14 Sep 1986, Ixtapantango, México, Mexico (ny 3902334).

**Figure 16.**
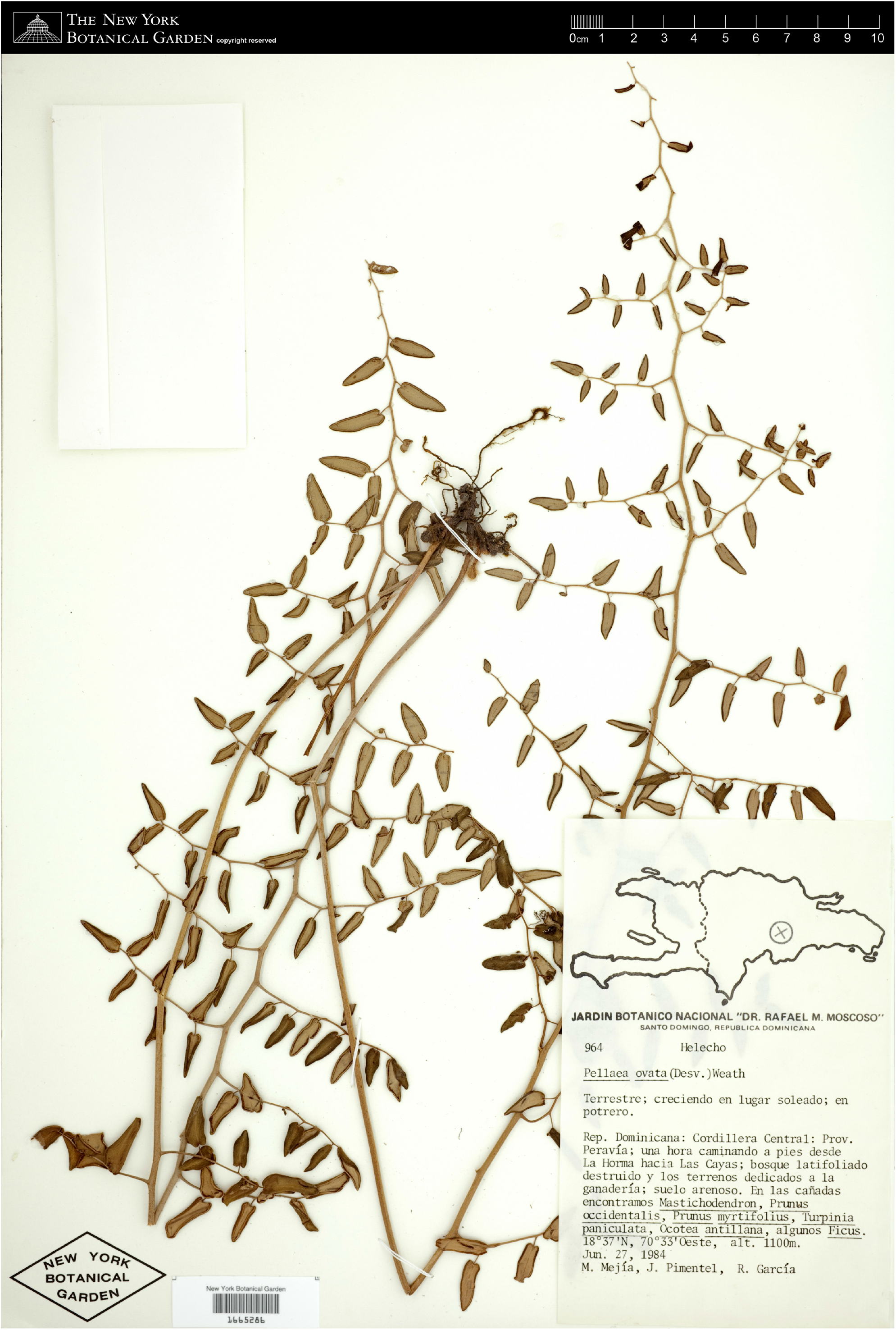
*Mejía &al. 964*, 27 Jun 1984, between La Horma &Las Cayas, Peravía, Dominican Republic (ny 1665286).

*Rachides* straight basally, weakly flexuous to nearly straight distally, glabrous; *costae* weakly flexuous, slightly reflexed to divaricate at base, usually arcing toward the frond apex, sometimes (especially in more basal pinnae) stiffly spreading at right angles to the rachis or slightly reflexed, glabrous; *stalks* of the pinnules (1-)2-5(−10) mm, weakly reflexed or, especially near the terminal pinnules, at right angles to the costae, glabrous; *fertile pinnules* narrowly ovate to broadly lanceolate, 2-3(−4) times longer than wide, *base* truncate to broadly and shallowly cordate, occasionally more deeply incised but for little more than the width of the stalk, usually oblique on terminal pinnules, sometimes oblique throughout, *sori* extending to the rounded-acute apex or nearly so.

*Pellaea ovata* s.s. overlaps these plants in most features, although the Hispaniola plant have rachides &costae consistently near the straight extreme, and pinnules near the narrow extreme, of *Pellaea ovata*. So far as I can tell given the limitations of specimen images, the Hispaniola plants have rachides &costae entirely glabrous, even the stalks of the pinnules glabrous. The rachides of *Pellaea ovata* can be puberulent only distally, or glabrescent with age, but there do not appear to be any without at least the stalks of the pinnules and distal third of the costae clearly puberulent. The pseudo-dichotomous pinnae &truncate pinnule apices of *Pellaea zygophylla* make it more obviously distinct from the plants on Hispaniola.

Dominican Republic. Azua: *García 2429* (ny 1665284). Independencia: *Zanoni 26434* (huap 27834, mexu 1403971); *Zanoni 37897* (ny 1665287). La Vega: *Abbott 21027* (UC 1871869); *Mejia 8840* (huap 27834, mexu 1403971); *Tuerckheim 2914* (ny 1665288); *Zanoni &Mejia 20786* (us 3257433); *Zanoni 17415* (ny 1665290). Peravia: *Mejia &al. 964* (huap 27834, mexu 1403971).

Haiti. Ouest: *Leonard 4804* (huap 27834, mexu 1403971).

### Pellaea sagittata

Although Mickel &Beitel (1988) believed *Pellaea oaxacana* to be a hybrid between *Pellaea ovata* and *Pellaea sagittata*, the distinction between *Pelleaa oaxacana* and *Pellaea sagittata* is clear. In addition to the characters mentioned in the keys of Mickel &Beitel (1988) and Mickel &Smith (2004), *Pellaea sagittata* has stipes that are sparsely scaly to the base of the blade. The rachides &often the costae are very sparsely scaly as well. The scales are relatively dense and conspicuous on leaves that are still unfurling. Although they are somewhat deciduous, at least a few scales persist well up the stipe or rachis in older leaves. The rhizome scales, also, are tan to rusty and concolorous in *Pellaea sagittata*, bicolorous and dark reddish brown to black centrally in *Pellaea oaxacana*. So long as mature leaves are present and their characters can be adequately observed, I have found no specimens or observations that are ambiguous and cannot be assigned to one species or the other.

However, in reviewing photographs of specimens and live plants, it became apparent that *Pellaea sagittata* is heterogeneous. Most photographs on iNaturalist show plants that are mostly or completely glabrous, but a few are conspicuously puberulent on the rachides &costae, and pubescent on the pinnules as well, especially towards the margins, sometimes across both adaxial and abaxial surfaces. Tryon (1957) highlights pubescence as a feature of *Pellaea sagittata* (as *Pellaea sagittata* var. *sagittata*), indicating that the “rachis and segment stalks [are] usually puberulous.” As mentioned above, she further remarks that pubescence, especially on the rachis, marks this apogamous taxon as well as apogamous plants of *Pellaea ovata*. Mickel &Smith (2004), on the other hand, state that the leaves of *Pellaea sagittata* are “glabrous or rarely sparsely puberulous”.

There is also considerable variation in other features. The leaves may be stiffly erect, with strongly ascending pinnae, V-shaped in cross section with pinnules folded upwards on each side of the costae; or spreading, the pinnae more weakly ascending &plane. The pinnules may be widely hastate &about as long as wide, to lanceolate and, in the most extreme plants, 4-5 times longer than wide. The stalks of the pinnules may be 1-2 mm long, with the cordate bases of the pinnules overlapping the costae, or 4-7 mm long, giving the leaves an open appearance. The veins are typically distinct &apparent on both surfaces of the pinnules but can be obscure adaxially or, less often, obscure on both surfaces. The puberulent plants are also toward the erect-leaved, short-stalked, narrow-pinnuled, and indistinct-veined end of the spectrum, and seem to be most frequent in the Mexican states of Chiapas, México, Michoacán, Oaxaca, and Puebla. Plants in South America, on the other hand, are generally toward the other end of the spectrum: glabrous, long-stalked, with widely hastate pinnules, and distinctly veined. Some of the South American specimens have both leaves with pinnules like those of *Pellaea cordifolia* and leaves with the pinnules much smaller, widely hastate, and often curled. The *Pellaea cordifolia-*like leaves appear to be produced earlier in the season, the leaves with hastate pinnules later. The extreme forms are distinctive but the variation between the extremes is extensive and complicated. It is not clear if there are taxa hiding within the mess. Some puberulent herbarium specimens are cited below, and specimens of the puberulent form and typical South American form are shown in Figures 18 &19.

## Pubescent specimens

Mexico. Chiapas: *Breedlove 40463* (ny 3902441); *Breedlove 51942* (ny 3902451). Chihuahua: *Knobloch 5983* (us 1791244). Ciudad de MÉxico: *Schaffner 90* (ny 3902457). MÉxico: *Hubert s*.*n*. (uc 2017350); *Matuda &al. 26719* (us 2083839); *Rose &Painter 7041* (us 450612); *Rose &Painter 7853* (us 451469); *Schumann 1900* (us 828038; rightmost leaf); *Tejero 2528* (ny 3902455). MichoacÁn: *Arsène 3629* (us 1030117); *Arsène 9986* (us 100016); *Feddema 31* (ny 3902442). Oaxaca: *Mickel 1648* (ny 3902463); *Smith 2058* (us 312921). Puebla: *Arsène 9970* (us 1030147). Querétaro: *Aguilar 68* (ny 3902452). San Luis PotosÍ: *Schaffner s*.*n*. (ny 3902464).

## Acknowledgements

This work would not be possible without data provided by many botanists, professional &amateur. This includes 234 contributors to iNaturalist in Mexico, Texas, Ecuador, &Peru who have photographed live *Pellaea ovata* s.l., and the many botanists and herbarium staff who have collected *Pellaea ovata* s.l., mounted and databased specimens, and taken photographs of specimens. I used images and data from the following herbaria:

Herbarium Berolinense (b) at Botanischer Garten und Botanisches Museum, Berlin.

BRIT Herbarium (brit) at Botanical Research Institute of Texas, Ft. Worth.

Desert Botanical Garden Herbarium (des), Phoenix.

John G. Searle Herbarium (f) at Field Museum of Natural History, Chicago.

Herbarium (hal) of Martin Luther University Halle-Wittenberg.

Howard Payne University Herbarium (hpc), Brownwood. Herbario Jardín Botánico Universitario (huap), Puebla.

Indiana University Herbarium (ind), Bloomington.

Herbario Real Jardín Botánico (ma), Madrid.

Herbario Nacional (mexu) at Universidad Nacional Autónoma de México, Mexico City.

University of Michigan Herbarium (mich), Ann Arbor. Mississippi State University Herbarium (missa), Starkville. Missouri Botanical Garden Herbarium (mo), St. Louis.

Michigan State University Herbarium (msc), East Lansing.

University of North Carolina at Chapel Hill Herbarium (ncu), Chapel Hill.

William and Lynda Steere Herbarium (ny) at the New York Botanical Garden, New York.

Vascular plants (p) at Muséum national d’Histoire naturelle, Paris.

Herbarium (ph) at Academy of Natural Sciences of Drexel University, Philadelphia.

Herbarium (rsa) at California Botanic Garden, Claremont.

Angelo State Natural History Collections Herbarium (sat), San Angelo.

University of Tennessee Herbarium (tenn), Knoxville.

Nationaal Herbarium Nederland (u) at Naturalis Biodiversity Center, Leiden.

University Herbarium (uc) at University of California, Berkeley.

United States National Herbarium (us) at Smithsonian Institution, Washington.

Garrett Herbarium (ut) at Utah Museum of Natural History, Salt Lake City.

Pringle Herbarium (vt) at University of Vermont, Burlington.

Wisconsin State Herbarium (wis) at University of Wisconsin, Madison.

Burke Museum Herbarium (wtu) at University of Washington, Seattle.

Yale University Herbarium (yu) at Yale University, New Haven.

Online specimen data portals are invaluable &greatly appreciated. I used data provided by the participants of the Consortium of California Herbaria (https://ucjeps.berkeley.edu/consortium/), the PteridoPortal network (https://www.pteridoportal.org/portal/), and SEINet (https://swbiodiversity.org/seinet/). These portals use the Symbiota content management system (https://symbiota.org), which is unequalled in this role thanks in large part to the tireless efforts of Ed Gilbert. Matt von Konrat and Daniel Le of the Field Museum kindly sent high resolution images of several specimens. Chris Hoess provided helpful discussion &pointed me towards some material I might otherwise have overlooked. Carl Rothfels provided comments on a draft that improved the clarity of the manuscript.

This article presents the understanding of the author, who is not acting as a representative of the Bureau of Land Management.

---

**Map 1.**
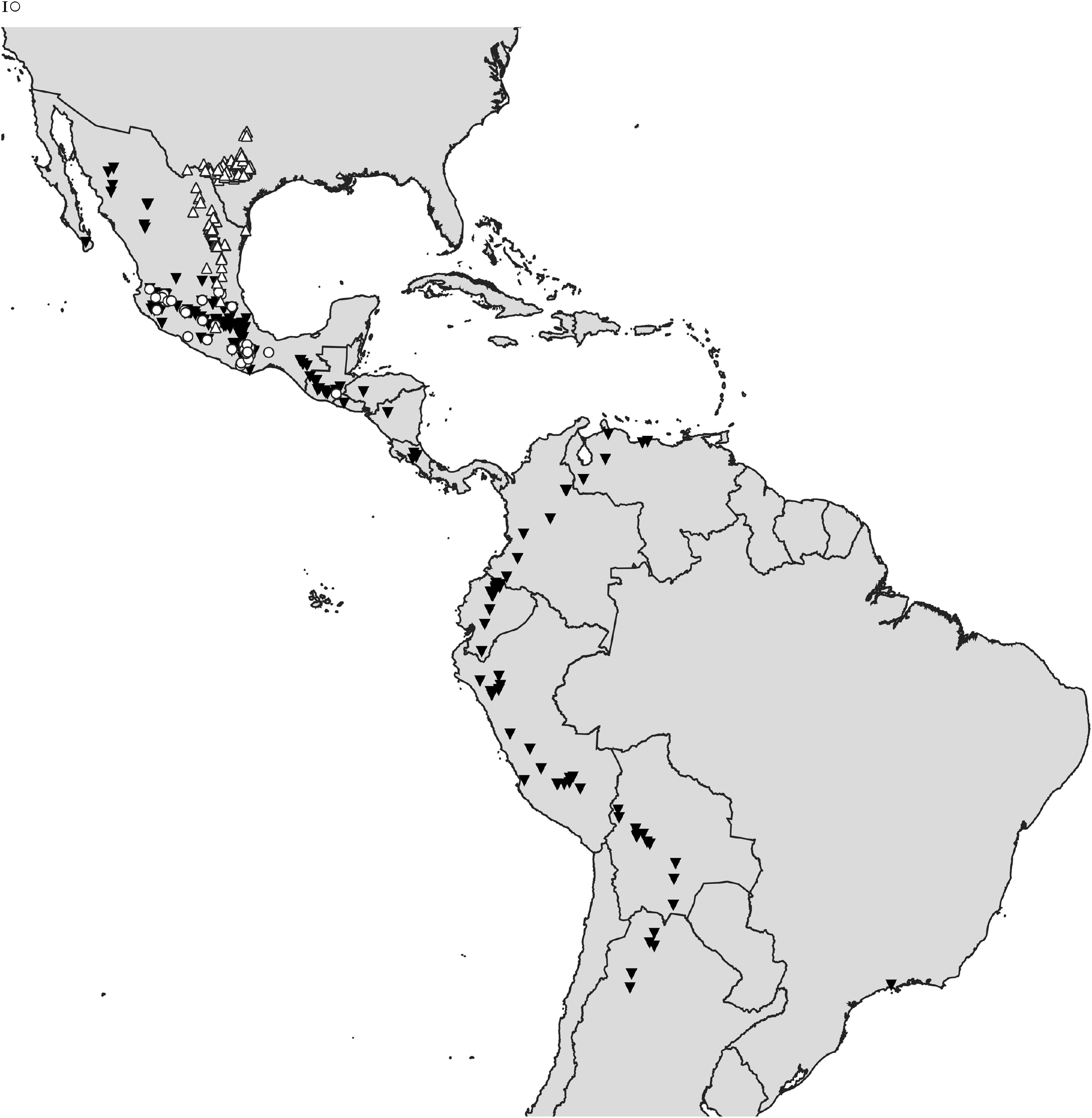
Complete geographic distribution of *Pellaea zygophylla*△,*Pellaea ovata*▾,and *Pellaea oaxacana*○.

**Map 2.**
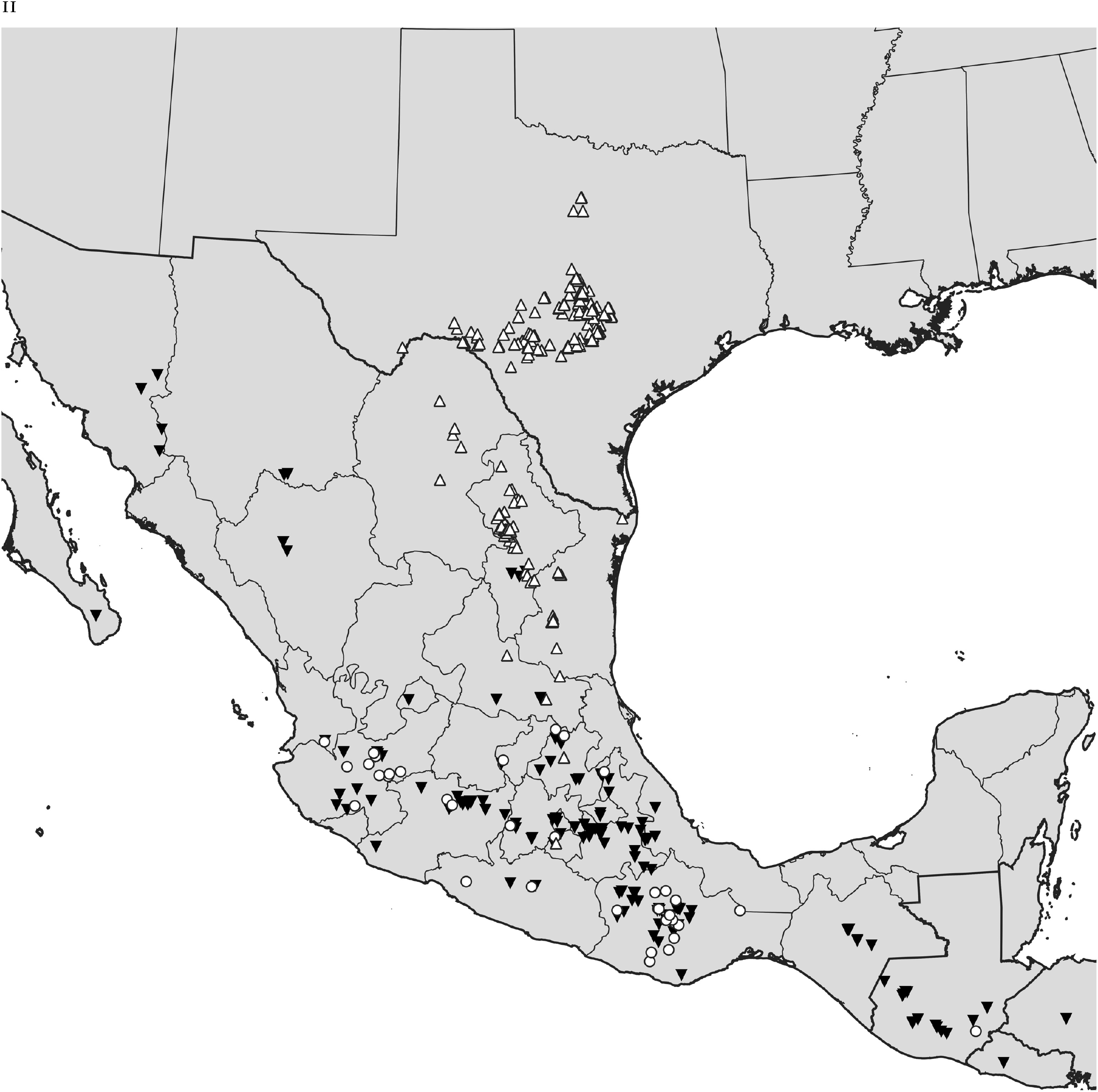
Geographic distribution of *Pellaea zygophylla*△,*Pellaea ovata*▾,and *Pellaea oaxacana*○,in Texas, Mexico, & Guatemala.

Following pages, Figures 2-6, *Pellaea zygophylla*.

Following pages, Figures 7-13, *Pellaea ovata*.

Following pages, Figures 14 &15, *Pellaea oaxacana*.

Following pages, Figures 16 &17, *Pellaea ovata* s.l. from Hispaniola.

Following pages, Figures 18 &19, *Pellaea sagittata*.

**Figure 17.**
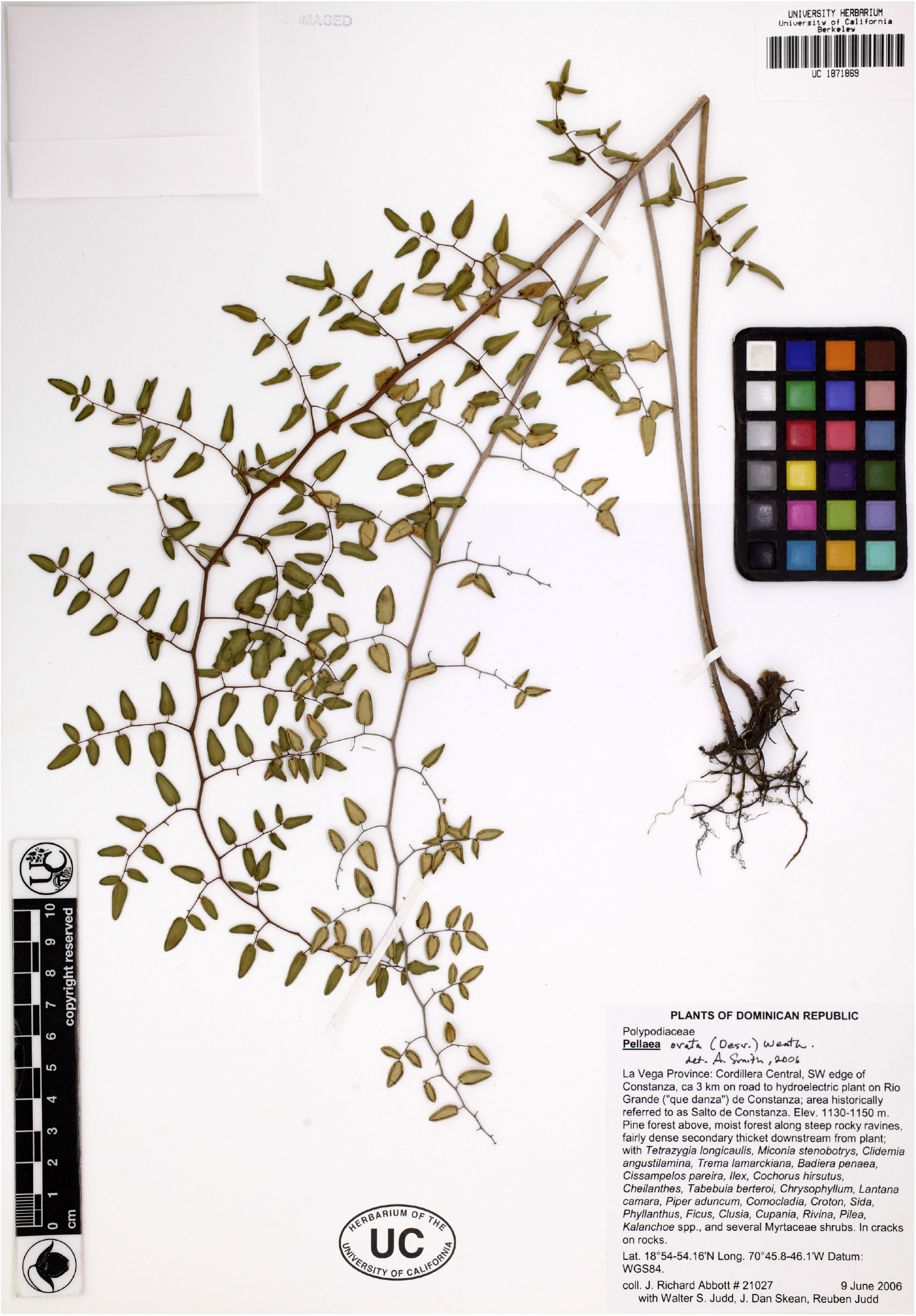
*Abbott &al. 21027*, 9 Jun 2006, Constanza, La Vega, Dominican Republic (UC 1871869).

**Figure 18.**
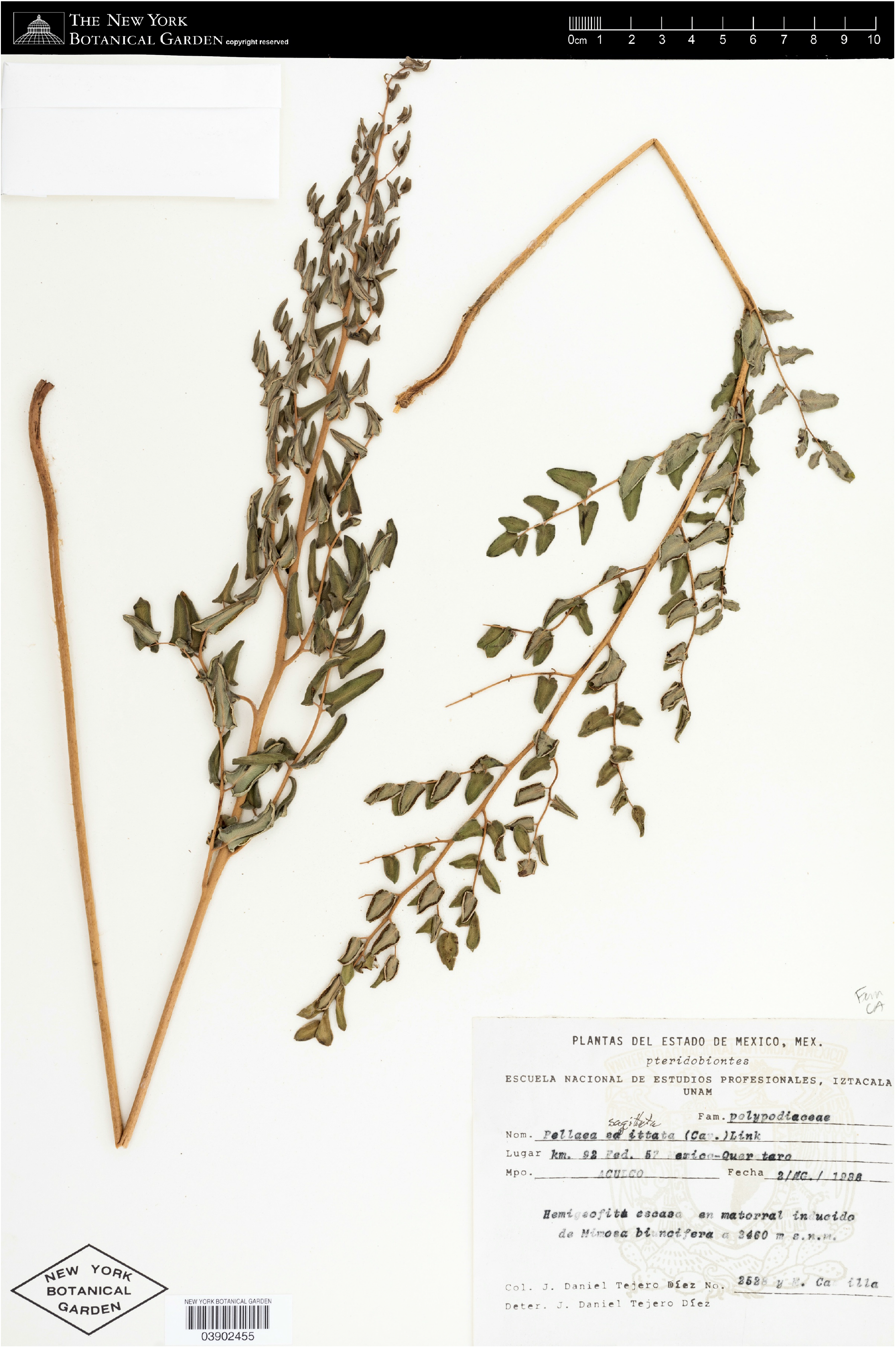
A puberulent specimen, *Tejero 2528*, 2 Aug 1986, between Ciudad de Querétaro &Ciudad de México, México, Mexico (NY 3902455).

**Figure 19.**
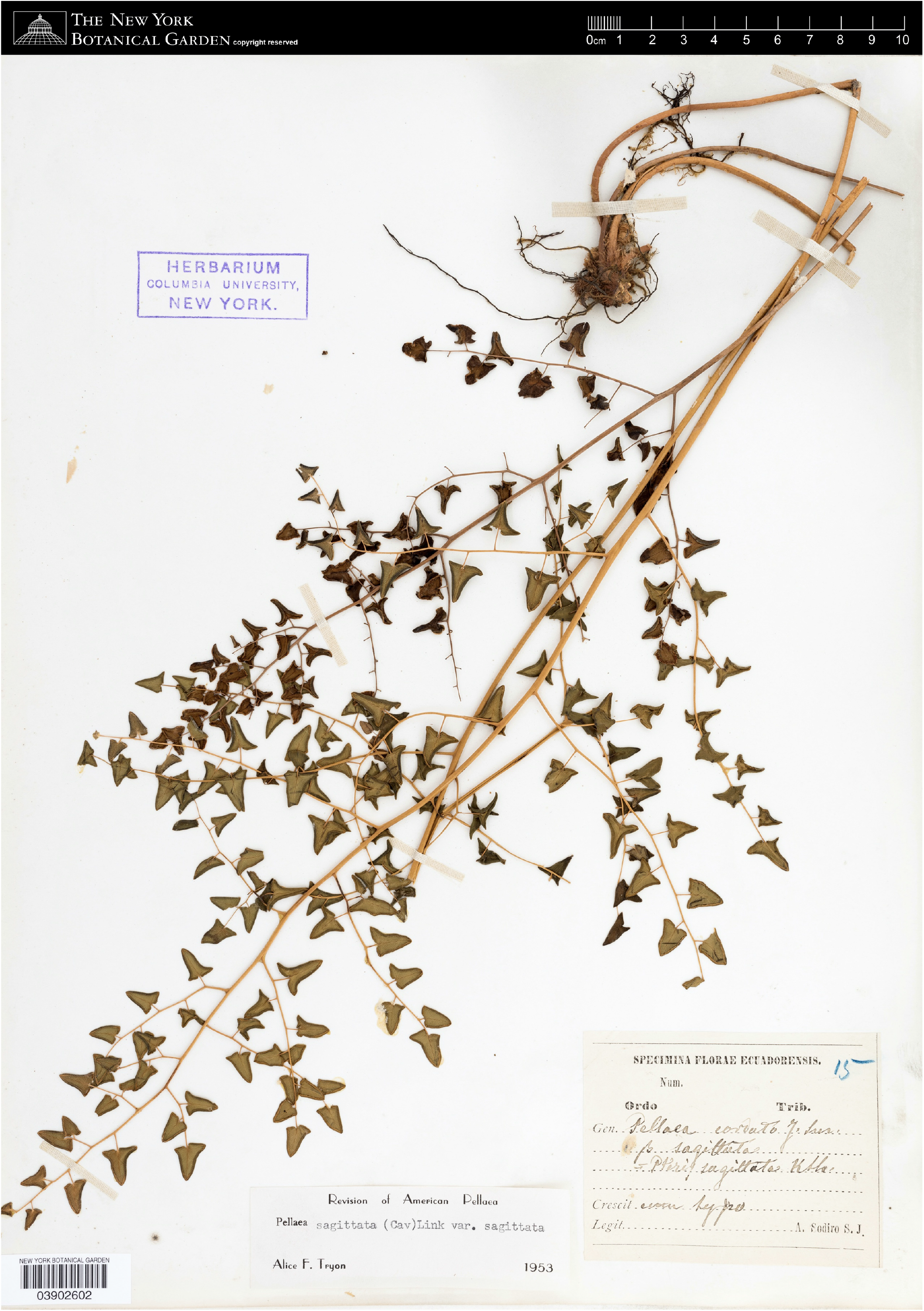
The typical South American form, *Sodiro 15*, s.d., Ecuador (NY 3902602).

